## Supplemental Information for "Structure in motion: visual motion perception as online hierarchical inference"

**6 Captions to the supplemental videos** **28**

### 1. Supplemental figures

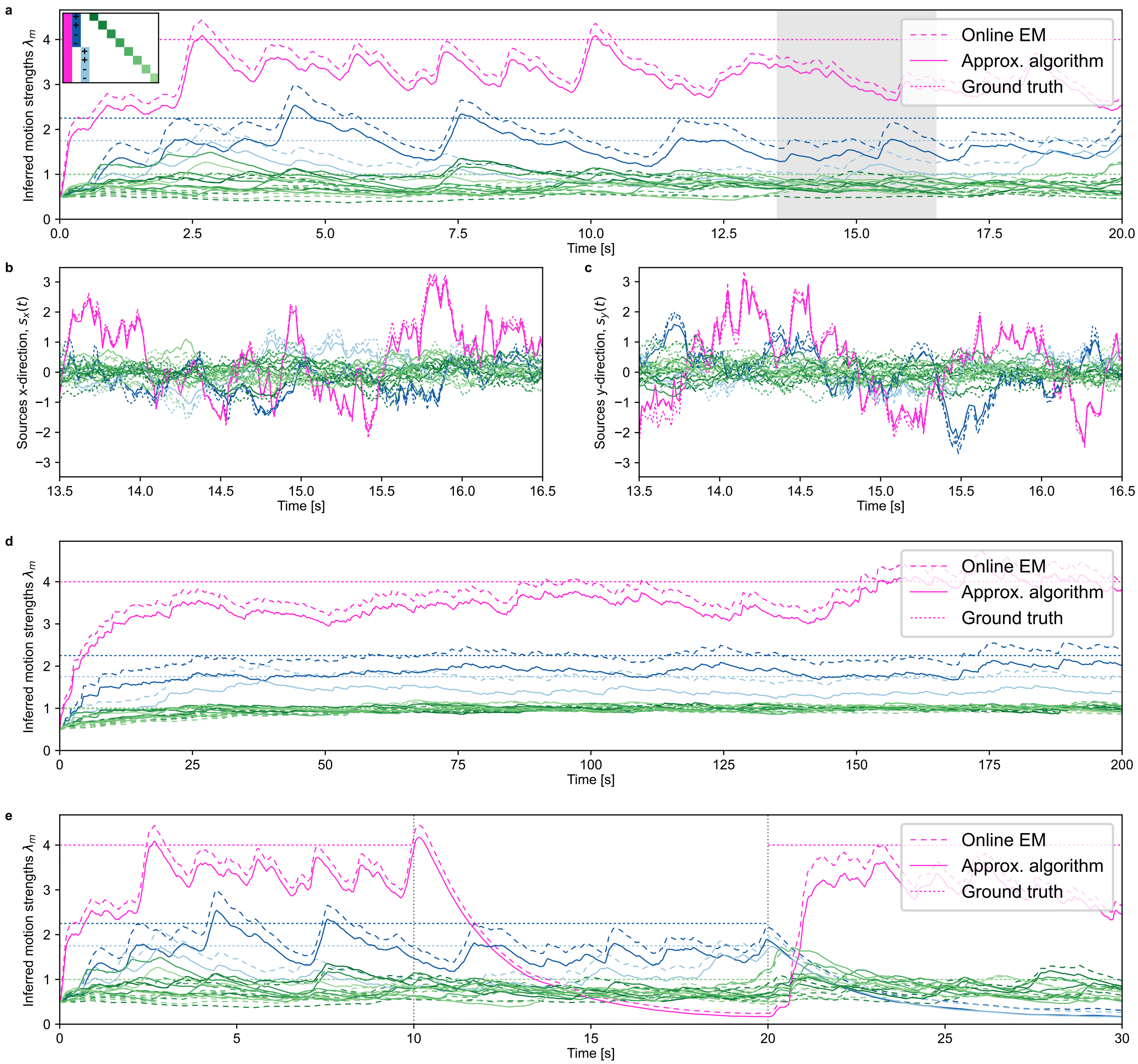

**Figure S1. The online inference model correctly recovers the structure and motion sources of presented input. (a)** Inferred motion strengths by the model. Two-dimensional input was generated from the generative model for a deeply nested structure with shared motion (pink; $\lambda=4$), two separate groups of counter-rotating sub-groups (dark- and light-blue; $+/-$ in the inset indicates $C_{km} = +1/-1$; $\lambda$=2.25 and 1.75), and eight individual motions (greens; $\lambda=1$). Other parameters are the default parameters for object-indexed experiments (see table in the *Methods* section of the main paper). Shown are the inferred strengths for the online inference model (solid lines; labeled "Approx. algorithm", given by eqn. (1)$-$(3) of the main text), the more accurate, but computationally also more complex online EM algorithm (dashed lines; given by eqn. (29), (30), and (35) of the Supplemental Information), and the ground truth (dotted lines). The approximate algorithm yields results similar to the reference online EM algorithm. Both algorithms underestimate the motion strengths due to the sparsity prior $p(\boldsymbol{\lambda}^2)$. **(b)** Inferred motion sources, x-direction for the highlighted duration of the simulation in panel a. Same color key as in panel a. **(c)** Same as panel b, but for the y-direction. **(d)** Repetition of the simulation in panel a, but with 10x longer time constant $\tau_\lambda$, longer run time, and uniform prior over the motion strengths. The underestimation in the reference algorithm vanishes; the approximate algorithm maintains its approximation quality. **(e)** Repetition of the simulation in panel a, but with a temporally changing structure. After 10 s, the shared component is switched off in the input. After 20 s, the shared component is re-introduced, but the groups are switched off. Both inference algorithms successfully detect these changes.

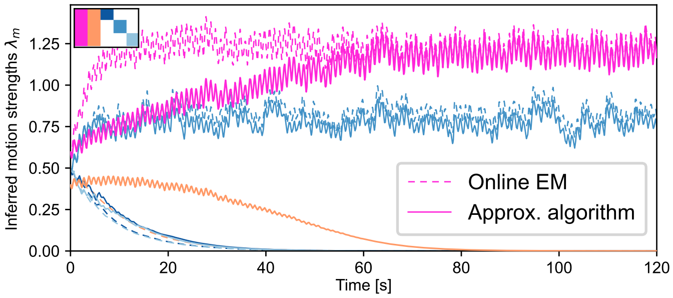

**Figure S2. The online model prefers simple structures, recruiting only necessary components from the reservoir.** Shown is a repetition of the Johansson experiment from Figure 2c, yet with a duplicated shared motion component in the observer model (pink and orange, see inset in the top-left). A small difference at initialization ($t=0$) between the two components widens, such that eventually only one component is recruited and the other one is dismissed. This preference for simpler structures is a direct consequence of the sparsity-inducing Jeffreys prior. If a uniform prior had been used, both shared components would have been maintained (not shown). Furthermore, we notice that the reference online EM algorithm converges more rapidly than the approximate algorithm. The reason is found in the posterior covariance matrix, $\mathbf{\Sigma}$, which is fully computed for online EM according to eqn. (29) and in which the off-diagonal element between the two shared components introduces competition during the credit assignment in eqn. (39) (uncertainty in the two sources is negatively correlated, leading to a negative matrix element in $\mathbf{\Sigma}$).

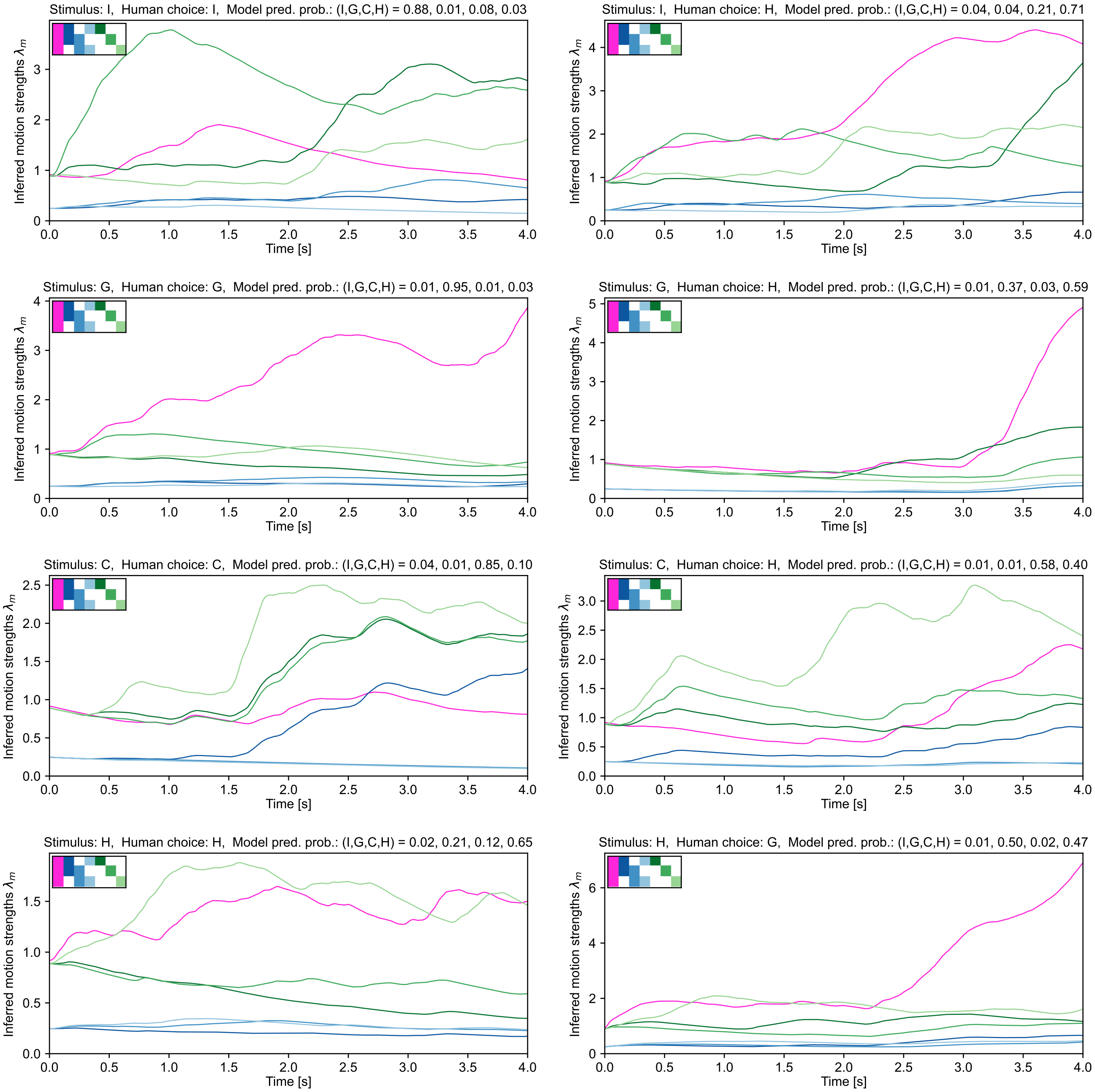

**Figure S3. Examples of motion structure inference for trials from (Yang et al., 2021).** Shown are traces for $\lambda(t)$ for eight example trials of participant #1. Axes titles state the ground truth, the participant's classification, and the predicted choice probabilities of the model. *Left column:* Trials of each structure which were correctly classified by the human participant. *Right column:* Trials of each structure which were incorrectly classified by the human participant.

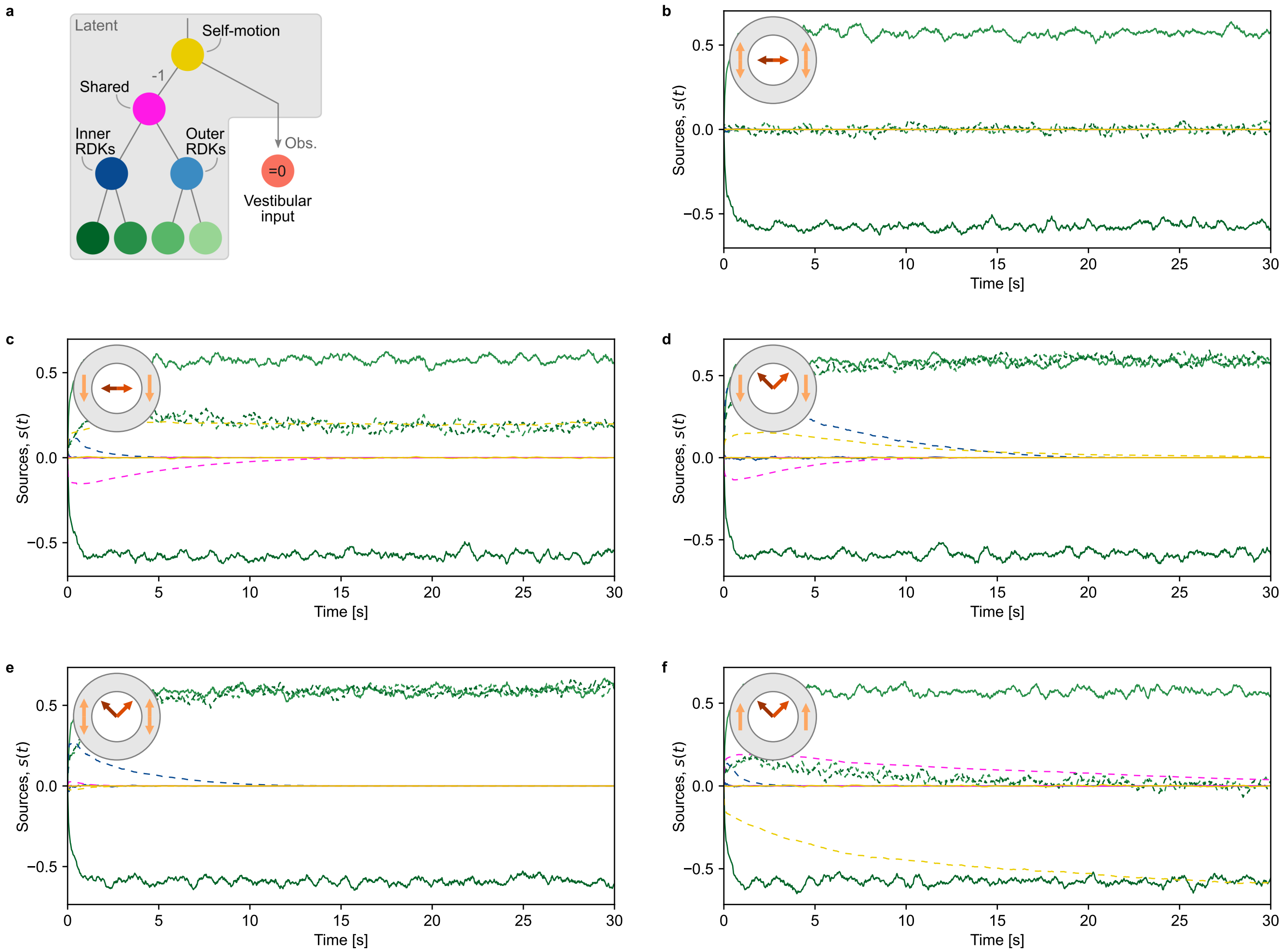

**Figure S5. Inferred motion sources for the center-surround interaction MDR experiment by (Takemura et al., 2011).** (a) Used motion tree with four hierarchy levels. (b)–(f) Example trials for each of the experiment conditions from Figure 4h–l of the main text. Shown is the evolution of motion sources (solid lines: x-direction, dashed lines: y-direction, colors as in panel (a)). For visual clarity, only the "inner branch" (left half of the tree) is shown, and traces were smoothed with a 500 ms box filter for plotting (the histograms in Figure 4 are based on the non-smoothed data). The perceived directions in Figure 4 are the sum of the $s_m$'s at trial end, without self-motion.

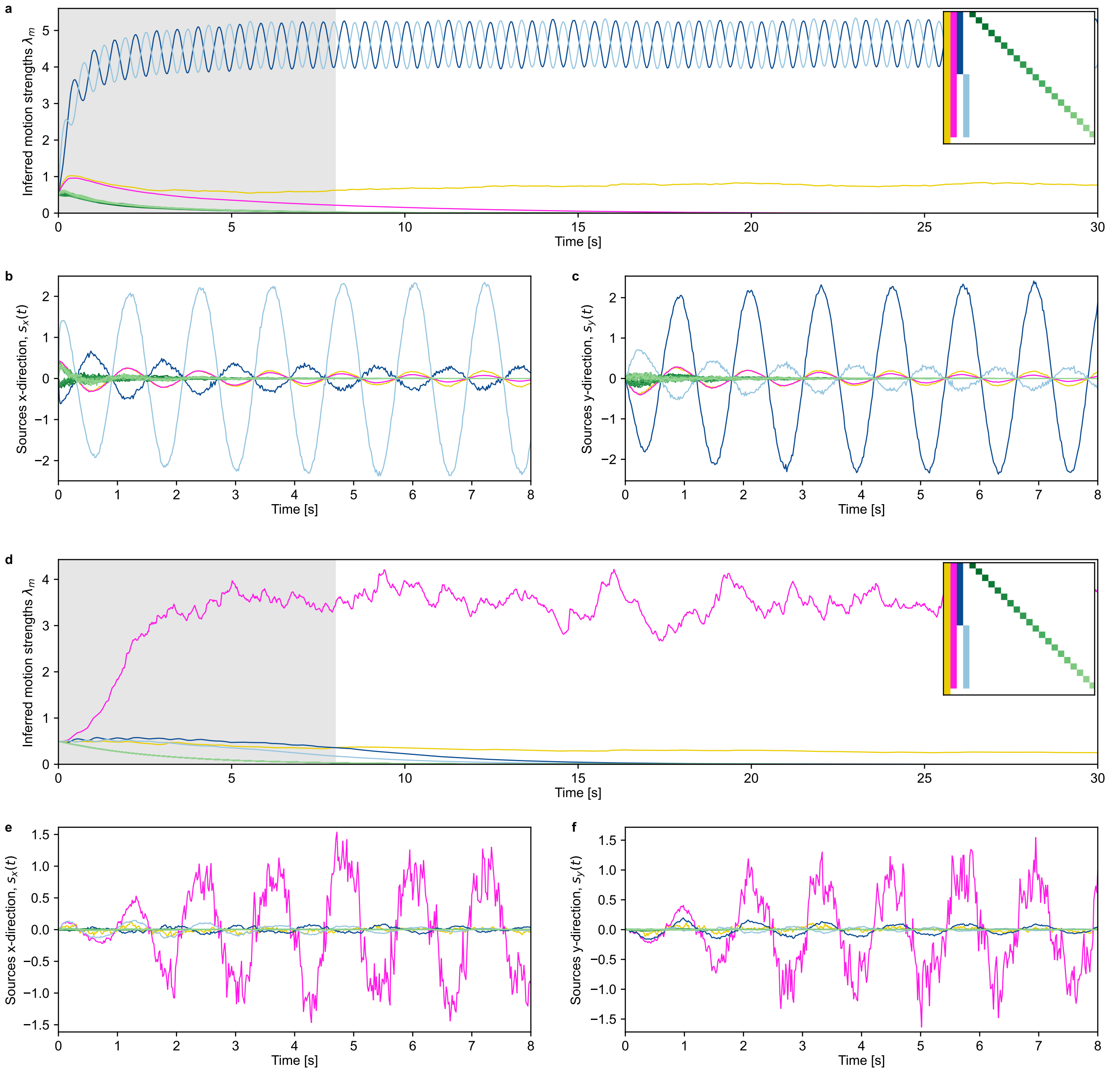

**Figure S6. Detailed evolution of all motion strengths and sources in the illusion from Lorenceau. (a)** Motion strengths, $\lambda(t)$, in the low-noise condition. Motion is decomposed into a deep hierarchy of self-motion (yellow), shared motion (magenta), group motion (dark and light blue), and individual motion of all $K=20$ dots (greens). In the generative model, self-motion affects the velocities of all dots and of the zero-mean vestibular input (lowest row in the $C$-matrix inset). **(b)** Inferred x-direction, $\mu_x(t)$, of all motion sources in the low-noise condition, for the highlighted duration in panel a. **(c)** Same as panel b, but for the y-direction. **(d–f)** Same as panels a–c, but for the high-noise condition.

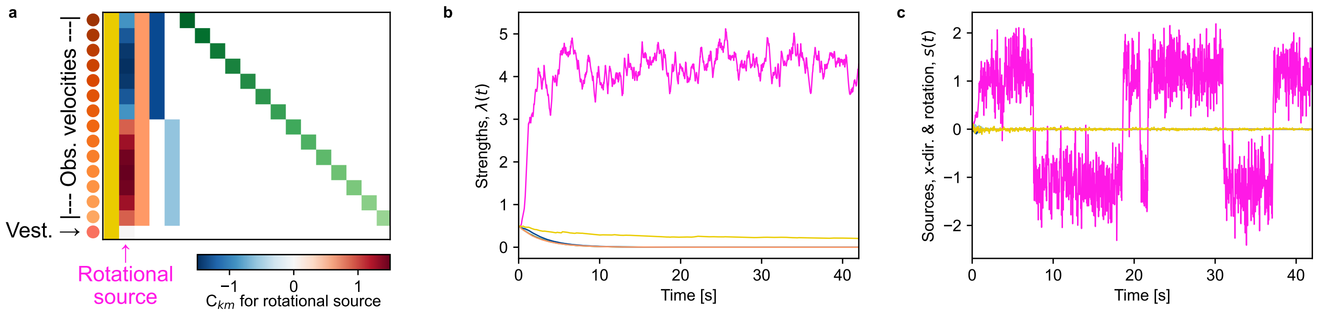

**Figure S7. A rotating cylinder is perceived for SfM also when more motion components are available. (a)** Component matrix with additional motion components. From left to right: Translational self-motion, rotational motion around vertical axis, translational motion for all visual inputs, translational motion for all dots on the back of the cylinder, translational motion for all dots on the front of the cylinder, translational motions for each individual observed velocity. All translational motions have $\|C_{km}\| = 1$ or $0$, and their colors indicate the line colors in panels (b) and (c). For the rotational motion component, the color map shows the value for the $v_x$-direction, i.e., $v_{x,k}(t) = C_{k,2}\, s^{\text{rot}}(t) +$ contributions of other sources. The locations of the observed velocities lie on the cylinder starting on the right and then ascending in CCW direction when viewed from the top (cf. Figure 6b of the main text). Thus, $s^{\text{rot}} > 0$, i.e., CCW rotation, leads to $v_x < 0$ for locations at the back of the cylinder. The rotational source, $s^{\text{rot}}$, is shown in magenta in panels (b) and (c) **(b)** Identified motion strengths. Even with the richer set of available components, the model identifies only the rotation (magenta). **(c)** Inferred motion sources. The bi-stability of the perceived rotation remains unaffected by the richer available structure (cf. Figure 6d of the main text; same random seed used for observation noise and stochastic time points of the assignment process).

#### List of used variables

| Variable | Description | Variable | Description |
|---|---|---|---|
| $\boldsymbol{v} = (v_1, .., v_K)^\top$ | Observable velocity | $\boldsymbol{s} = (s_1, .., s_M)^\top$ | Latent motion source |
| $v_k = v_{k,d}(t)$ | Dim. $d$ and time $t$ often suppressed | $s_m = s_{m,d}(t)$ | Dim. $d$ and time $t$ often suppressed |
| $\hat{\boldsymbol{v}} = (\hat{v}_1, .., \hat{v}_K)^\top$ | Noise-free velocity | $\boldsymbol{\lambda} = (\lambda_1, .., \lambda_M)^\top$ | Motion strength |
| $\boldsymbol{\mu} = (\mu_1, .., \mu_M)^\top$ | Mean vector in $\boldsymbol{s}$-posterior | $\boldsymbol{\epsilon} = (\epsilon_1, .., \epsilon_K)^\top$ | Prediction error |
| $\boldsymbol{\Sigma} = \boldsymbol{\Omega}^{-1}$ | Covariance in $\boldsymbol{s}$-post. and inv. precision | $f_\Sigma$ | Adiabatic approx. of $\sigma^2$ |
| $\boldsymbol{\sigma}^2$ | Vect. of diag. elements $(\Sigma_{11}, .., \Sigma_{MM})^\top$ | $\boldsymbol{C}$ | Component matrix of shape $(K \times M)$ |
| $D$ | no. dimensions | $\boldsymbol{c}_m$ | m-th column of $\boldsymbol{C}$ |
| $\mathrm{diag}[\boldsymbol{x}]$ | Diag. matrix over some vec. $\boldsymbol{x}$ | $\langle f(x) \rangle_{p(x)}$ | Expectation of $f(x)$ under $p(x)$ |
| $\tau_s$ | Time const. for $s$-inference (OU process) | $\boldsymbol{J}$ | Interaction prior on motion comp. |
| $\tau_\lambda$ | Time const. for $\lambda$-inference | $\delta t$ | Obs. interval (inverse frame rate) |
| $\tau_\epsilon$ | Time const. for pred. err. | $\delta(t)$, $\delta_{ij}$ | Dirac and Kronecker delta |
| $t_{\mathrm{obs}}$ | Obs. time point in cont.-time (CT) | $\partial_t$ | Partial derivative (here, w.r.t. $t$) |
| $t$ | No. time steps (DT); <u>Or:</u> time (CT) | $\boldsymbol{\nu}$ | No. of $\boldsymbol{\mu}$-pseudo obs. (hyperprior) |
| $\mathcal{I}\chi(\nu, \kappa^2)$ | Scaled inverse chi-squared distribution | $\boldsymbol{\kappa}$ | Val. of pseudo observations |
| $\sigma_\rho$ | MT width of speed tuning | $\kappa_\alpha$ | MT precision of direction tuning |
| $\sigma_{\mathrm{obs}}$ | Observation noise ($\delta t$-independent) | $\mathcal{Q}(\boldsymbol{\lambda})$ | EM expect. compl. data log-likelihood |
| $\widetilde{\sigma}_{\mathrm{obs}}$ | Disc.-time obs. noise ($=\sigma_{\mathrm{obs}} / \sqrt{\delta t}$) | $S$ | Structure $I, G, C, H$ in (Yang et al., 2021) |
| $R$ | Radial rec. field loc. | $\beta$ | Inv. temp. in (Yang et al., 2021) |
| $\vartheta$ | Angular rec. field location | $b_G, b_C, b_H$ | Biases in (Yang et al., 2021) |
| $s_r, s_\varphi$ | Polar sources: radial and angular velocity | $\pi_{\mathrm{L}}$ | Lapse prob. in (Yang et al., 2021) |
| $\rho$ | Speed in MT tuning func. | $\alpha$ | Direction (angle) in MT tuning |
| $n_\rho$, $n_\alpha$, $N_\rho$, $N_\alpha$ | MT neuron indices, max. values | $\mu_\rho(n_\rho)$, $\mu_\alpha(n_\alpha)$ | Tuning center of neuron $(n_\rho, n_\alpha)$ |
| $\psi$ | MT max. firing rate multiplier | $I_n(\kappa)$ | Modified Bessel function of order $n$ |
| $f, f_\sigma, f_\rho, f_\alpha$ | MT tuning func. and sub-functions | $\boldsymbol{I}$ | Identity matrix |
| $\boldsymbol{W}, \overline{\boldsymbol{W}}$ | Linear maps in algo. & neural domain | $\boldsymbol{Q}, \overline{\boldsymbol{Q}}$ | Quad. maps in algo. & neural domain |
| $\boldsymbol{b}, \overline{\boldsymbol{b}}$ | Add. constants in algo. & neural domain | $\boldsymbol{A}, \boldsymbol{A}^\dagger$ | Linear decoding matrix & adjoint |
| $\boldsymbol{x}, \boldsymbol{y}, \boldsymbol{z}$ | Variables in generic network derivation | $x_k$, $y_k$ | Spatial locations for 2-dim. stimuli |
| $\boldsymbol{r}_{\mathrm{inp}}$, $\boldsymbol{r}_{\mathrm{dis}}$, $\boldsymbol{r}_{\text{1-to-1}}$ | Firing rates in the network model | $\gamma$ | Direction repulsion opening angle |
| $v_{\mathrm{vst}}$ | Vestibular self-motion input | $\dot{\varphi}$ | Angular (rotational) velocity |
| $\sigma_{\mathrm{vst}}$ | Obs. noise for vestibular input | $\dot{r}$ | Radial velocity |
| $\eta_C$ | Learning rate for $\boldsymbol{C}$-learning | | |

**Table 1. List of used variables.** In a few cases, we chose to accept notation clashes when confusion is ruled out and the dual use actually facilitates clarity. These cases are indicated with “<u>Or:</u>” in the description.

#### Theory of the online hierarchical inference model and the neural network model

In **Section 2**, we introduce the generative model for structured motion. In **Section 3**, we derive the online hierarchical inference model. In **Section 4**, we present optional extensions to the inference model. In **Section 5**, we derive the recurrent neural network model.

#### 2. Generative model of structured motion

The following model of hierarchically structured motion is an adaptation of the generative model from (Bill et al., 2020). We consider $K$ observable velocities, $v_{k,d}(t)$, in $D$ spatial dimensions. To prevent clutter, we will develop most of the theory for the one-dimensional case, $D{=}1$, and use the vector notation, $\boldsymbol{v}{=}(v_1,..,v_K)^\top$. The extension to $D{>}1$ is straightforward and will be covered in a dedicated subsection. We will often suppress the explicit time dependence when confusion is ruled out. In other cases, we may write time as an index, $\boldsymbol{v}_t$, when a compact notion is desirable. Observable velocities, $\boldsymbol{v}$, are noisy instantiations of noise-free velocities, $\hat{\boldsymbol{v}}$, which are generated by $M$ latent motion sources, $s_{m,d}(t)$, as will be specified below. Similar to velocities, we abbreviate $\boldsymbol{s}{=}(s_1,..,s_M)^\top$.

##### 2.1 Composition of observable velocity from motion motifs

For the purpose of this work, we restrict the influence of motion source $s_m(t)$ on velocity $\hat{v}_k(t)$ to three particularly basic relations: the motion of $s_m$ can affect $\hat{v}_k$ either positively (e.g., the latent flock motion on an observed bird's velocity), negatively (e.g., the effect of self-motion on the observed scene), or not at all (e.g., the flock motion's effect on a tree). Formally, we describe these influences in a $K \times M$ component matrix $\boldsymbol{C}$, with $C_{km} = +1$, $-1$, and 0 for positive, negative and absent influence, respectively. We will sometimes refer to the columns of $\boldsymbol{C}$ as motion motif or motion component $\boldsymbol{c}_m$. Typically, we have $M > K$ because every observable can have its individual motion component which exclusively affects this observable.

The overall velocity $\hat{v}_k$ is the sum of all the motion sources' contributions:

$$\hat{\boldsymbol{v}} = \boldsymbol{C}\,\boldsymbol{s} \ . \tag{1}$$

It is easy to see that all tree-like hierarchies can be cast into this form. As a word of warning, the opposite is not true: not every possible $\boldsymbol{C}$ represents hierarchically structured motion.

Finally, it is worth mentioning that motion components are not necessarily limited to the simple $C_{km} = \pm 1$ values. For modeling gradual relationships, a motion source could affect some observables stronger than others. For instance, the current of a river could influence the water's velocity in the middle of the river stronger than close to the bank. All subsequent derivations likewise hold for such gradual relationships, as long as they influence the observed velocities linearly.

##### 2.2 Generation of observable velocities from stochastic, latent motion sources

For the generative model, we assume that the motion sources, $s_m(t)$, evolve independently according to an Ornstein-Uhlenbeck process, see, e.g., (Gardiner, 2009),

$$\mathrm{d}s_m = -\tfrac{1}{\tau_s} s_m \,\mathrm{d}t + \lambda_m \,\mathrm{d}W_m \ , \tag{2}$$

where Wiener process $W_m(t)$ drives changes in $s_m$ via random forces, motion strength $\lambda_m \geq 0$ controls the magnitude of these forces, and $\tau_s > 0$ is the typical time constant of significant changes in $s_m$. The equilibrium distribution of this process is a normal distribution with zero mean and variance $\frac{\tau_s}{2}\lambda_m^2$:

$$\lim_{t\to\infty} p(s_m(t)) = \mathcal{N}\left(s_m; 0, \tfrac{\tau_s}{2}\lambda_m^2\right) \ . \tag{3}$$

The Ornstein-Uhlenbeck process is a suitable first-order approximation for modeling real-world motion as it (i) generates continuous trajectories $s_m(t)$, (ii) includes a notion of inertia/momentum via its temporal filtering with $\tau_s$, (iii) offers an intuitive parameterization by scaling typical velocities linearly in $\lambda_m$, and (iv) implements a slow-velocity prior via a normal distribution like in (Weiss, Simoncelli, and Adelson, 2002).

Observable velocities, $\boldsymbol{v}(t)$, are generated by composing the latent motion sources to noise-free velocities according to eqn. (1) and then applying independent Gaussian noise:

$$\boldsymbol{v}(t) \sim \mathcal{N}\left(\boldsymbol{C}\,\boldsymbol{s}(t), \widetilde{\sigma}_{\mathrm{obs}}^2 \boldsymbol{I}\right) \ . \tag{4}$$

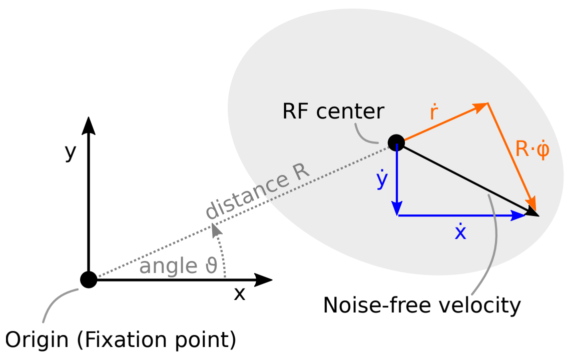

$$\dot{x} = \dot{r}\,\cos\vartheta - \dot{\varphi}\,R\sin\vartheta$$
$$\dot{y} = \dot{r}\,\sin\vartheta + \dot{\varphi}\,R\cos\vartheta$$

$$\dot{r} = \dot{x}\,\cos\vartheta + \dot{y}\,\sin\vartheta$$
$$\dot{\varphi} = \frac{1}{R}\left(-\dot{x}\,\sin\vartheta + \dot{y}\,\cos\vartheta\right)$$

**Figure S8. Coordinate transformation from polar motion sources to Cartesian velocities for supporting rotational and radial motion motifs.** When receptive field (RF) centers are fixed, rotational motion, $\dot{\varphi}$, and radial motion, $\dot{r}$, feature fixed linear relations to the resulting Cartesian velocities in x- and y-direction. The angles $\vartheta$ and $\varphi$ are measured relative to the x-axis in counter-clockwise direction by convention. Cartesian components are measured in rightward (x) and upward (y) direction.

In foresight of the continuous-time formulation, we denote the observation noise by $\widetilde{\sigma}_{\text{obs}}$ (with a tilde). It will later be adjusted to the frame rate of incoming observations such that the information per unit time remains constant. To keep the derivation tractable, we will ignore the reported velocity-dependence of observation noise in human perception (Weber's law) and treat $\widetilde{\sigma}_{\text{obs}}$ as a constant.

Marginalizing over the stationary distribution of the latent motion sources, eqn. (3), the observable velocities are jointly normally distributed due to their linear dependence on a Gaussian origin, with zero mean and covariance matrix $\frac{\tau_s}{2}\boldsymbol{C}\,\text{diag}\left[\boldsymbol{\lambda}^2\right]\boldsymbol{C}^\top + \widetilde{\sigma}^2_{\text{obs}}\,\boldsymbol{I}$, with $\text{diag}\left[\boldsymbol{\lambda}^2\right]$ denoting the diagonal matrix generated from vector $\boldsymbol{\lambda}^2 = (\lambda_1^2, .., \lambda_M^2)^\top$.

#### 2.3 Motion structure

The motion strengths $\boldsymbol{\lambda}$ play a particularly important role in the generative process described by eqn. (2)+(4). For $\lambda_m = 0$, dependent motion sources decay to zero, i.e., $s_m \to 0$. Hence, motion strengths describe the presence ($\lambda_m > 0$) or absence ($\lambda_m = 0$) of motion components, as well as their typical magnitude ($\langle |s_m| \rangle \propto \lambda_m$). In other words, given a reservoir of components, $\boldsymbol{C}$, which have been learned to occur in visual scenes in general, the vector $\boldsymbol{\lambda}$ will describe the structural composition of motion relations in a specific visual scene. Knowing $\boldsymbol{\lambda}$ therefore is equivalent to knowing the motion structure of the scene.

#### 2.4 Extension to multiple spatial dimensions

For $D > 1$, we will usually assume that all $s_{m,d}$ with the same index $m$ share motion strength $\lambda_m$ and motion component $\boldsymbol{c}_m$, yet each $s_{m,d}$ follows its own stochastic evolution, i.e., has its own Wiener process, $W_{m,d}$, in eqn. (2). This choice reflects that space is isotropic: due to the Gaussianity of each $s_{m,d}$, the joint distribution $p(s_{m,1}, .., s_{m,D})$ will be a multivariate Gaussian with covariance matrix $\frac{\tau_s}{2}\lambda_m^2\,\boldsymbol{I}$, with $\boldsymbol{I}$ denoting the identity matrix, and is, thus, invariant to rotations in the experimenter's choice of the coordinate system. Again, this is a useful first-order approximation, even though human motion perception has been reported to be not perfectly isotropic (Manning, Thomas, and Braddick, 2018; Moscatelli et al., 2019).

In summary, the generative model of structured motion is characterized by the following set of parameters: number of spatial dimensions $D$, motion strengths $\boldsymbol{\lambda}$, component matrix $\boldsymbol{C}$, time constant $\tau_s$, and observation noise $\widetilde{\sigma}_{\text{obs}}$. Some extensions of this model covering heterogeneous time constants, non-isotropic observation noise and missing observations will be discussed in dedicated subsections in the context of online inference, below.

#### 2.5 Polar coordinates: rotational and radial motion

The discussion so far assumed that motion sources affect velocities in a translational (Cartesian) manner, that is, by adding the same vector to all dependent velocities. An important exception are rotational and radial motion in two dimensions ($D=2$) which typically occur in flow field parsing (when moving forward, everything expands on the retina; when tilting your head, everything rotates in the opposite rotational direction).

It turns out that rotational and radial motion sources can be incorporated into our framework of linear velocity generation as per eqn. (1), when the spatial locations of all dependent velocities are fixed. This is, for example, fulfilled in experimental setups using drifting gratings: the drifting grating has motion energy, but its location stays fixed at the same spot. Note how this setup is different from the perspective commonly taken in physics where velocity entails changes in location. The perspective we take here is geared towards studying brain computation where a group of neurons often processes motion in a certain, fixed area of the inputs. Specifically, this is satisfied in many visual areas including V1, MT, and MST, where neurons feature a fixed spatial receptive field in retinal coordinates. The index $k$ in $v_k$ then refers to a fixed location on the retina. We therefore call this condition "location-indexed".

To derive the generative model for rotational and radial motion in a location-indexed experiment, consider the spatial receptive fields illustrated in **Fig. S8**. Each receptive field (RF) has a fixed center relative to the fovea and is characterized by radial distance $R$ and angle $\vartheta$, which is measured relative to the x-axis in counter-clockwise direction by convention. The generative model can use latent motion sources in radial direction, denoted by $\dot{r}$ in the figure, and in angular direction, denoted by $\dot{\varphi}$. These polar motion sources generate velocities $\dot{x}$ and $\dot{y}$ in Cartesian space. Importantly, the (generative) transformation from polar to Cartesian coordinates is linear for each receptive field:

$$\begin{aligned} \dot{x} &= \dot{r}\cos\vartheta - \dot{\varphi}\,R\sin\vartheta \\ \dot{y} &= \dot{r}\sin\vartheta + \dot{\varphi}\,R\cos\vartheta \ . \end{aligned} \tag{5}$$

Consequently, the generative process can be included into the component matrix $\boldsymbol{C}$ when each polar motion source affects dependent observables, $\hat{\boldsymbol{v}}_k$, in both spatial dimensions. In contrast to the previously discussed translational motion sources, rotational and radial motion sources maintain separate motion components $\boldsymbol{c}_m$ and motion strengths $\lambda_m$:

$$\begin{aligned} c_{k,m=\mathrm{rad},d=x} &= \cos\vartheta_k \ , & c_{k,m=\mathrm{rot},d=x} &= -R_k\sin\vartheta_k \\ c_{k,m=\mathrm{rad},d=y} &= \sin\vartheta_k \ , & c_{k,m=\mathrm{rot},d=y} &= +R_k\cos\vartheta_k \ . \end{aligned} \tag{6}$$

As before, the model adds up the velocities from different sources and applies the observation noise only to the final velocity (in Cartesian space). The example of rotational and radial motion demonstrates how spatial dimensions can be mixed, as long as the coordinate transformation is linear for every observable $\hat{\boldsymbol{v}}_k$.

**Rotation in 3D and structure-from-motion.** The above addresses rotations around a pivot point on the fovea, i.e., the rotation happens in the same plane as the observations. A different form of rotational motion, which is equally covered by the generative model, is used for the structure-from-motion experiments in Figure 6 of the main text. Here, the rotation is around an axis that lies in the observation plane (e.g., the y-axis in the x-y-plane) while the motion happens in 3D, including a depth component, $\hat{v}_{k,z}$. As illustrated in Figure 6b of the main text, rotation (and also expansion around the axis) lead to a linear relation between the rotational motion source, $s_t^{\mathrm{rot}}$, and the noise-free velocity vectors, $\hat{\boldsymbol{v}}_t$, at every fixed location in location-indexed experiments. As before, noise is added in Cartesian coordinates. Note that in the SfM displays of the main text, no depth information is presented in order to make the stimuli ambiguous with regard to their direction of rotation.

#### 3. Online hierarchical inference

In **Section 3.1**, we develop the hierarchical inference algorithm in a discrete-time, batch formulation, using the Expectation-Maximization (EM) algorithm. In **Section 3.2**, we draw the continuous-time limit to obtain an online algorithm. Finally, in **Section 3.3**, we introduce an adiabatic approximation which reduces the required computations to neuron-friendly operations. This is the online model underlying all simulations of the main paper.

##### 3.1 Inference via the Expectation-Maximization algorithm

Our goal for motion structure inference is to simultaneously infer the value of motion sources $\boldsymbol{s}(t)$ and the underlying structure $\boldsymbol{\lambda}$ from a stream of observations $\boldsymbol{v}_{1:t} = (\boldsymbol{v}_1, .., \boldsymbol{v}_t)$ with observations arriving at time steps of duration $\delta t$ (inverse frame rate). The number of spatial dimensions $D$, components $\boldsymbol{C}$, time constant $\tau_s$, and observation noise $\widetilde{\sigma}_{\mathrm{obs}}$ are assumed to be known (although $\boldsymbol{C}$ could be learned on longer time scales, see **Section 4.5**). The challenge in this hierarchical inference task is that $\boldsymbol{s}_t$ and $\boldsymbol{\lambda}$ are mutually dependent on another: $\boldsymbol{\lambda}$ acts as a parameter in $p(\,\boldsymbol{s}_t\,|\,\boldsymbol{\lambda}\,)$ per eqn. (2), and will therefore affect the posterior $p(\,\boldsymbol{s}_t\,|\,\boldsymbol{v}_{1:t}\,)$. On the other hand, inferring the presence of, say, flocking birds ($\lambda_{\mathrm{flock}} > 0$) depends on "perceiving" an instantaneous flock motion, $s_{\mathrm{flock}}(t)$, in the first place.

The EM algorithm (Dempster, Laird, and Rubin, 1977) offers a solution to this chicken-and-egg problem. For its application, we leverage the fact that motion sources and strengths change on different time scales, $\tau_s$ for $\boldsymbol{s}_t$ and $\tau_\lambda$ for $\boldsymbol{\lambda}$. For $\tau_\lambda \gg \tau_s$, we can treat $\boldsymbol{\lambda}$ as a constant while inferring $\boldsymbol{s}(t)$—known as the E-step in EM—, and then, in alternation, optimize $\boldsymbol{\lambda}$ based on the inferred motion strengths—the M-step in EM. In **Section 3.1.1** and **Section 3.1.2**, we will address the E-step and the M-step separately. As before, we will develop the theory in one spatial dimension for notational clarity and then generalize to $D > 1$. Further, we will present the derivation in discrete time in a batch formulation. The continuous-time limit will be drawn in **Section 3.2**.

##### 3.1.1 Inference of motion sources for a given structure (E-step)

For the E-step, we aim to infer $p(\boldsymbol{s}_{1:t} \,|\, \boldsymbol{v}_{1:t}\,;\boldsymbol{\lambda})$ for a given structure $\boldsymbol{\lambda}$, and then compute the expected value of the log-likelihood of the augmented data distribution $p(\boldsymbol{v}_{1:t}, \boldsymbol{s}_{1:t}\,; \boldsymbol{\lambda})$, see, e.g., Section 9.3. in (Bishop, 2006). For the remainder of this subsection, we will often suppress the explicit dependence on $\boldsymbol{\lambda}$ to avoid notational clutter. Since we are interested in an online algorithm, we will use the filtering solution which is obtained from iterative application of temporal propagation to the next observation time,

$$p(\boldsymbol{s}_{t-1} \,|\, \boldsymbol{v}_{1:t-1}) \longrightarrow p(\boldsymbol{s}_t \,|\, \boldsymbol{v}_{1:t-1}) \ , \tag{7}$$

and integration of the next observation,

$$p(\boldsymbol{s}_t \,|\, \boldsymbol{v}_{1:t}) \propto p(\boldsymbol{s}_t \,|\, \boldsymbol{v}_{1:t-1})\, p(\boldsymbol{v}_t \,|\, \boldsymbol{s}_t) \ . \tag{8}$$

Propagation, eqn. (7), is performed by propagating the density according to the stochastic process in eqn. (2). Mathematically, this is done by convolving $p(\boldsymbol{s}_{t-1} \,|\, \boldsymbol{v}_{1:t-1})$ with the Gaussian transition density $p(\boldsymbol{s}_t \,|\, \boldsymbol{s}_{t-1})$ of the Ornstein-Uhlenbeck process. Integration, eqn. (8), is the application of Bayes rule using the emission model of eqn. (4).

For linear stochastic dynamics with a Gaussian emission model, i.e., the present case, the posterior will always be a multivariate Gaussian with some mean $\boldsymbol{\mu}_t$ and covariance $\boldsymbol{\Sigma}_t$: $p(\boldsymbol{s}_t \,|\, \boldsymbol{v}_{1:t}) = \mathcal{N}(\boldsymbol{s}_t; \boldsymbol{\mu}_t, \boldsymbol{\Sigma}_t)$. Kalman filtering (Kalman, 1960) is one possible algorithm for calculating the posterior moments $\boldsymbol{\mu}_t$ and $\boldsymbol{\Sigma}_t$, and we refer the interested reader to the Supporting Information of (Bill et al., 2020) for explicit forms of the Kalman filter's state transition matrix and process noise covariance matrix. For the present work, we will employ a more elegant, continuous-time solution that is equivalent to the Kalman-Bucy filter (Kalman and Bucy, 1961; Jazwinski, 2007) for calculating $\boldsymbol{\mu}_t$ and $\boldsymbol{\Sigma}_t$. The derivation, which is provided in **Section 3.2**, will furthermore facilitate a neuro-friendly approximate implementation (**Section 3.3**). For now, let us assume that the posterior moments, $\boldsymbol{\mu}_{1:t}$ and $\boldsymbol{\Sigma}_{1:t}$, have been computed by whichever method of choice.

**E-step.** We compute the expected value of the log-likelihood of the augmented data distribution,[1]

$$Q(\boldsymbol{\lambda}) = \langle \log p(\boldsymbol{v}_{1:t}, \boldsymbol{s}_{1:t}\,; \boldsymbol{\lambda}) \rangle_{p(\boldsymbol{s}_{1:t} \,|\, \boldsymbol{v}_{1:t})} \tag{9}$$

$$= \sum_{j=1}^{t} \langle \log p(\boldsymbol{v}_j \,|\, \boldsymbol{s}_j\,; \boldsymbol{\lambda}) + \log p(\boldsymbol{s}_j \,|\, \boldsymbol{s}_{j-1}\,; \boldsymbol{\lambda}) \rangle_{p(\boldsymbol{s}_j, \boldsymbol{s}_{j-1} \,|\, \boldsymbol{v}_{1:t})} \tag{10}$$

$$\overset{\text{ind.}}{\approx} \sum_{j=1}^{t} \langle \log p(\boldsymbol{v}_j \,|\, \boldsymbol{s}_j\,; \boldsymbol{\lambda}) + \log p(\boldsymbol{s}_j\,; \boldsymbol{\lambda}) \rangle_{p(\boldsymbol{s}_j \,|\, \boldsymbol{v}_{1:j})} \overset{\text{def.}}{=} \sum_{j=1}^{t} Q_j(\boldsymbol{\lambda}) \ , \tag{11}$$

where, at the third equality, we have made two approximations, namely, that (i) consecutive observations were independent, and (ii) we take the expectation w.r.t. the filtering posterior, $p(\boldsymbol{s}_j \,|\, \boldsymbol{v}_{1:j})$, rather than the smoothing density, $p(\boldsymbol{s}_j \,|\, \boldsymbol{v}_{1:t})$. The first approximation will be corrected for during the M-step in **Section 3.1.2**, where we will weigh the likelihood term as if it was comprised of only $t \cdot \delta t/\tau_s$ independent samples when combining it with a sparsity prior on $\boldsymbol{\lambda}$. This correction is justified because it is equivalent to considering only a sparse subsample of observations that each lie $\tau_s$ apart. Consecutive observations in this subsample are almost decoupled since the Ornstein-Uhlenbeck process in eqn. (2) decorrelates the motions sources at time scale $\tau_s$. The second approximation—filtering rather than smoothing—is a modeling assumption for what information is employed by an online agent, e.g., a human observer.

[1]In the 2nd line, we implicitly assume, for mathematical rigor only, that there is an initial distribution, $p(\boldsymbol{s}_{t=0})$, which is absorbed again in the 3rd line.

The $Q_j(\boldsymbol{\lambda})$ can be calculated analytically:

$$Q_j(\boldsymbol{\lambda}) = \langle \log p\left(\boldsymbol{v}_j \,|\, \boldsymbol{s}_j\,;\boldsymbol{\lambda}\right) + \log p(\boldsymbol{s}_j\,;\boldsymbol{\lambda})\rangle_{p\left(\boldsymbol{s}_j\,|\,\boldsymbol{v}_{1:j}\right)} \qquad \leftarrow \text{with } p(\boldsymbol{s}_j\,|\,\boldsymbol{v}_{1:j}) = \mathcal{N}\left(\boldsymbol{s}_j; \boldsymbol{\mu}_j, \boldsymbol{\Sigma}_j\right) \tag{12}$$

$$= \langle -\tfrac{1}{2}(\boldsymbol{v}_j - \boldsymbol{C}\boldsymbol{s}_j)^\top \tilde{\sigma}_{\text{obs}}^{-2}\boldsymbol{I}\,(\boldsymbol{v}_j - \boldsymbol{C}\boldsymbol{s}_j) - \tfrac{1}{2}\boldsymbol{s}_j^\top \text{diag}\big[\tfrac{\tau_s}{2}\boldsymbol{\lambda}^2\big]^{-1}\boldsymbol{s}_j - \tfrac{1}{2}\log|\tilde{\sigma}_{\text{obs}}^2\boldsymbol{I}| - \tfrac{1}{2}\log|\text{diag}\big[\tfrac{\tau_s}{2}\boldsymbol{\lambda}^2\big]\,|\rangle_{p\left(\boldsymbol{s}_j\,|\,\boldsymbol{v}_{1:j}\right)} + \text{const.} \tag{13}$$

$$= \sum_{m=1}^{M} -\frac{1}{\tau_s}\frac{\mu_{j,m}^2 + \sigma_{j,m}^2}{\lambda_m^2} - \frac{1}{2}\log\lambda_m^2 \quad [\text{dropping } \boldsymbol{\lambda}\text{-independent terms and using } \langle s^2\rangle_{\mathcal{N}(s;\mu,\sigma^2)} = \mu^2 + \sigma^2] \tag{14}$$

with $\sigma_{j,m}^2 := \Sigma_{j,mm}$, $|\cdot|$ denoting the determinant, and the $\lambda$-independent terms have been dropped because they will not play a role in the maximization w.r.t. $\boldsymbol{\lambda}$ in **Section 3.1.2**. For the full $Q(\boldsymbol{\lambda})$, we thus obtain:

$$Q(\boldsymbol{\lambda}) = \sum_{j=1}^{t} Q_j(\boldsymbol{\lambda}) = -\frac{t}{\tau_s}\sum_{m=1}^{M}\frac{\langle \mu_{j,m}^2 + \sigma_{j,m}^2\rangle_j}{\lambda_m^2} + \frac{\tau_s}{2}\log\lambda_m^2\ , \tag{15}$$

with $\langle\cdot\rangle_j$ being the time average over the batch.

##### 3.1.2 Inference of motion strengths and sparsity prior (M-step)

For the M-step, $Q(\boldsymbol{\lambda})$ is maximized w.r.t. $\boldsymbol{\lambda}$ to obtain the maximum likelihood (ML) solution. Here, we will make use of the additional freedom to impose a prior distribution, $p(\boldsymbol{\lambda})$, on the motion strengths, see, e.g., Section 9.4. in (Bishop, 2006). First, we will introduce a family of prior distributions which reflect our knowledge that from a reservoir of motion components most components will be absent or small in any given scene (sparsity prior). Then, we will perform the M-step to derive the maximum a posteriori (MAP) solution.

**Sparsity prior.** In foresight of the M-step, we formulate the prior over $\boldsymbol{\lambda}^2$ (instead of $\boldsymbol{\lambda}$) and choose

$$p(\boldsymbol{\lambda}^2; \boldsymbol{\nu}, \boldsymbol{\kappa}^2) = \prod_{m=1}^{M}\mathcal{I}\chi(\lambda_m^2; \nu_m, \kappa_m^2) \qquad \text{with} \qquad \mathcal{I}\chi(\lambda^2; \nu, \kappa^2) = \frac{1}{\lambda^{(2+\nu)}}\exp\left[-\frac{\nu\kappa^2}{2\lambda^2} - A(\nu,\kappa^2)\right]\ . \tag{16}$$

$\mathcal{I}\chi(\lambda^2;\nu,\kappa^2)$ denotes the density of the scaled inverse chi-squared distribution which is the conjugate prior to a normal distribution with known mean and unknown variance. Conceptually, this is exactly our task at hand: we know that $\langle s_m\rangle = 0$ in the generative model, but we have to estimate its variance, $\langle s_m^2\rangle$, which is controlled by $\lambda_m^2$. As will become obvious in the M-step, the (hyper-)parameters, $\boldsymbol{\nu}$ and $\boldsymbol{\kappa}^2$, will take the role of pseudo-counts and pseudo-observations, respectively. The log-partition, $A(\nu,\kappa^2) = \log\Gamma(\frac{\nu}{2}) - \frac{\nu}{2}\log\frac{\nu\kappa^2}{2}$, will have no effect on the MAP estimate.

Two choices of $\nu$ and $\kappa^2$ are of particular interest. For $\nu = \kappa^2 = 0$, we have $p(\lambda^2) \propto 1/\lambda^2$ which is a non-informative (Jeffreys) prior on the variance of $s^2$. For $\nu = -2$, $\kappa^2 = 0$, we have $p(\lambda^2) \propto 1$ which is a uniform prior. The latter choice of hyper-parameters will turn the MAP estimate into the ML estimate.

**M-step.** To maximize $Q$, we find the roots of its $\lambda_m^2$-derivatives:

$$0 \stackrel{!}{=} \frac{\mathrm{d}}{\mathrm{d}\lambda_m^2}\left(Q(\boldsymbol{\lambda}) + \log p(\boldsymbol{\lambda}^2; \boldsymbol{\nu}, \boldsymbol{\kappa}^2)\right) = \frac{t}{\tau_s}\left(\frac{\langle\mu_{j,m}^2 + \sigma_{j,m}^2\rangle_j}{(\lambda_m^2)^2} - \frac{\tau_s}{2}\frac{1}{\lambda_m^2}\right) + \frac{\nu_m\kappa_m^2}{2\,(\lambda_m^2)^2} - \frac{1+\frac{\nu_m}{2}}{\lambda_m^2} \tag{17}$$

$$\Rightarrow\ t\,\langle\mu_{j,m}^2 + \sigma_{j,m}^2\rangle_j + \tfrac{\tau_s}{2}\nu_m\kappa_m^2 = t\tfrac{\tau_s}{2}\lambda_m^2 + \tfrac{\tau_s}{2}(2+\nu_m)\lambda_m^2 \tag{18}$$

$$\Rightarrow\ \tfrac{\tau_s}{2}\lambda_m^2 = \frac{t\,\langle\mu_{j,m}^2 + \sigma_{j,m}^2\rangle_j + \frac{\tau_s}{2}\nu_m\kappa_m^2}{2+\nu_m+t}\ . \tag{19}$$

Eqn. (19) highlights several intuitive properties of motion structure inference. First, the MAP value of motion strength $\lambda_m^2$ only depends on the inferred posterior moments of the corresponding motion source $s_m$, that is, there is no cross-talk between motion sources $s_m$ and $s_{m'}$. Second, recalling that $\frac{\tau_s}{2}\lambda_m^2$ is the expected long-term variance of $s_m$ according to eqn. (3), eqn. (19) tells us to match this expected variance, $\frac{\tau_s}{2}\lambda_m^2$, to the observed variance, $\langle\mu_{j,m}^2 + \sigma_{j,m}^2\rangle_j$, of the inferred motion source *over time*.[2] Third, hyper-parameter $\kappa_m$ plays the role of an average pseudo-observation,

[2]This is to be distinguished from the posterior's instantaneous uncertainty $\sigma_{t,m}^2$.

$\frac{\tau_s}{2}\kappa_m^2 = \langle \mu_m^2 + \sigma_m^2 \rangle$, which is then weighted as $\nu_m$ pseudo-samples against the $t$ observed data samples. Thus, $\kappa_m$ describes (a priori) typical values of $\lambda_m$. (The summand 2 in the denominator of eqn. (19) is a relict of the scale-invariant Jeffreys prior.) Finally, a uniform hyper-prior, $\nu_m = -2$, $\kappa_m = 0$, yields straightforward variance matching as the ML solution.

We conclude the M-step, by correcting eqn. (19) for the fact that the $t$ data samples actually only represent $t \cdot \delta t / \tau_s$ independent samples, as promised in **Section 3.1.1**,

$$\frac{\tau_s}{2}\lambda_m^2 = \frac{\frac{t \cdot \delta t}{\tau_s} \langle \mu_{j,m}^2 + \sigma_{j,m}^2 \rangle_j + \frac{\tau_s}{2}\nu_m \kappa_m^2}{2 + \nu_m + \frac{t \cdot \delta t}{\tau_s}} \ . \tag{20}$$

Eqn. (20) is the batch solution to motion structure inference which we will build on for the continuous-time formulation in **Section 3.2**.

##### 3.1.3 Extension to multiple spatial dimensions

When $D > 1$, we typically assume that each $\lambda_m$ controls the variance of the $s_{m,d}$ in all spatial dimensions $d$ (see **Section 2.4**). When going through the derivation of the EM algorithm, the decisive changes happen in eqn. (15): (i) the expectation now runs over all spatial dimensions, i.e., $\langle \sum_{d=1}^{D} \mu_{j,m,d}^2 + \sigma_{j,m,d}^2 \rangle_j$; (ii) the log-partition gets multiplied by $D$, due to each $\lambda_m$ contributing with the power of $D$ to $\log|\mathrm{diag}[\boldsymbol{\lambda}^2]|$. With these changes, the M-step in eqn. (20) finds its optimum when

$$D\,\frac{\tau_s}{2}\lambda_m^2 = \frac{\frac{t \cdot \delta t}{\tau_s} \langle \sum_{d=1}^{D} \mu_{j,m,d}^2 + \sigma_{j,m,d}^2 \rangle_j + \frac{\tau_s}{2}\nu_m \kappa_m^2}{\frac{2}{D} + \nu_m + \frac{t \cdot \delta t}{\tau_s}} \ . \tag{21}$$

Here, we have made a slight re-parameterization of $\nu_m$ and $\kappa_m$ to preserve the developed intuition that $\kappa_m$ describes typical values of $\lambda_m$, and that $\nu_m$ counts the number of pseudo-observations.[3]

#### 3.2 Continuous-time, online inference

We now turn to a continuous-time formulation of the above motion stucture inference algorithm. While doing so, we will overload the notation of time $t$ which previously denoted integer-valued time steps and now becomes real-valued. We will point out the respective locations where this transition happens, below, to preclude confusion. Concretely, we will first reformulate the generative model in the form of natural parameters in **Section 3.2.1**. Then, we derive continuous-time dynamics on these parameters in **Section 3.2.2**, as had been promised in **Section 3.1.1**, for solving the E-step. Finally, we cast eqn. (21) into a recursive equation for an online, continuous-time M-Step in **Section 3.2.3**. This results in the reference *online EM algorithm* for online hierarchical motion structure inference.

##### 3.2.1 Equivalent formulation of the generative model and inference using natural parameters

Knowing that all distributions involved in propagation, eqn. (7), and integration, eqn. (8), are multivariate Gaussians, we write the emission model, eqn. (4), in terms of the sufficient statistics of $\boldsymbol{s}_t$,

$$p(\boldsymbol{v}_t \,|\, \boldsymbol{s}_t) = \mathcal{N}\left(\boldsymbol{v}_t; \boldsymbol{C}\boldsymbol{s}_t, \tilde{\sigma}_{\mathrm{obs}}^2 \boldsymbol{I}\right) \propto \exp\left[\begin{pmatrix} \boldsymbol{s}_t \\ \boldsymbol{s}_t\boldsymbol{s}_t^\top \end{pmatrix} \cdot \begin{pmatrix} \frac{\boldsymbol{C}^\top \boldsymbol{v}_t}{\tilde{\sigma}_{\mathrm{obs}}^2} \\ -\frac{1}{2}\frac{\boldsymbol{C}^\top \boldsymbol{C}}{\tilde{\sigma}_{\mathrm{obs}}^2} \end{pmatrix}\right] \ , \tag{22}$$

where we have dropped all $\boldsymbol{s}_t$-independent terms because they will play no role in the inference. Denoting the propagated, yet pre-integration, distribution by

$$p(\boldsymbol{s}_t \,|\, \boldsymbol{v}_{0:t-\delta t}) \propto \exp\left[\begin{pmatrix} \boldsymbol{s}_t \\ \boldsymbol{s}_t\boldsymbol{s}_t^\top \end{pmatrix} \cdot \begin{pmatrix} \boldsymbol{\Omega\mu}_t \\ -\frac{1}{2}\boldsymbol{\Omega}_t \end{pmatrix}\right] \ , \tag{23}$$

with yet-to-be-determined natural parameters $\boldsymbol{\Omega}_t := \boldsymbol{\Sigma}_t^{-1}$ and $\boldsymbol{\Omega\mu}_t := \boldsymbol{\Omega}_t \boldsymbol{\mu}_t$, the integration of observations, eqn. (8), amounts to the following simple updates:

$$\boldsymbol{\Omega}_t \mapsto \boldsymbol{\Omega}_t + \frac{\boldsymbol{C}^\top \boldsymbol{C}}{\tilde{\sigma}_{\mathrm{obs}}^2} \qquad \text{and} \qquad \boldsymbol{\Omega\mu}_t \mapsto \boldsymbol{\Omega\mu}_t + \frac{\boldsymbol{C}^\top \boldsymbol{v}_t}{\tilde{\sigma}_{\mathrm{obs}}^2} \ . \tag{24}$$

[3] Specifically, we substituted $\frac{\nu_m}{D} \to \nu_m$ and $D\kappa_m^2 \to \kappa_m^2$. Then, $\nu_m = 1$ means one pseudo-observation in *each* spatial dimension. A uniform prior is imposed by $\nu_m = -2/D$ and $\kappa_m = 0$.

In eqn. (23), we have made use already of the continuous-time notation, with time running from 0 to $t$ and observations arriving at $\delta t$-intervals. We next address propagation, i.e., the continuous-time dynamics of eqn. (2) in terms of $\boldsymbol{\Omega}\boldsymbol{\mu}_t$ and $\boldsymbol{\Omega}_t$.

##### 3.2.2 Continuous-time dynamics of natural parameters

For the OU process, eqn. (2), we know the evolution of the distribution of $\boldsymbol{s}_t$ between observations in closed form (see, e.g., (Gardiner, 2009)). Namely, the mean $\boldsymbol{\mu}_t$ decays towards $\mathbf{0}$ exponentially with time constant $\tau_s$, and the covariance $\boldsymbol{\Sigma}_t$ decays towards its steady state value $\mathrm{diag}\left[\frac{\tau_s}{2}\boldsymbol{\lambda}^2\right]$ exponentially with time constant $\frac{\tau_s}{2}$. From these known dynamics of $\boldsymbol{\mu}_t$ and $\boldsymbol{\Sigma}_t$, we calculate the dynamics of the natural parameters:

$$\partial_t \boldsymbol{\Omega}_t = -\boldsymbol{\Omega}_t \left(\partial_t \boldsymbol{\Sigma}_t\right) \boldsymbol{\Omega}_t = \ldots = \left(\tfrac{\tau_s}{2}\right)^{-1} \underbrace{\left(\boldsymbol{I} - \boldsymbol{\Omega}_t \,\mathrm{diag}\left[\tfrac{\tau_s}{2}\boldsymbol{\lambda}^2\right]\right)}_{\circledast} \boldsymbol{\Omega}_t + \frac{\boldsymbol{C}^\top \boldsymbol{C}}{\widetilde{\sigma}^2_{\mathrm{obs}}}\,\delta(t - t_{\mathrm{obs}}) \;. \tag{25}$$

For completeness, we have already included the integration of observations, eqn. (24), at observation time $t_{\mathrm{obs}}$ in the dynamics. In contrast to the dynamics of $\boldsymbol{\Sigma}_t$, the dynamics of $\boldsymbol{\Omega}_t$ are non-linear. Yet, we observe that, in the absence of observations, eqn. (25) leads to the desired fixed point since $\circledast = 0$ for $\boldsymbol{\Omega}^{-1} = \mathrm{diag}\left[\frac{\tau_s}{2}\boldsymbol{\lambda}^2\right]$. Likewise, we obtain for the other natural parameter $\boldsymbol{\Omega}\boldsymbol{\mu}_t$:

$$\partial_t (\boldsymbol{\Omega}\boldsymbol{\mu}_t) = (\partial_t \boldsymbol{\Omega}_t)\,\boldsymbol{\mu}_t + \boldsymbol{\Omega}_t\,(\partial_t \boldsymbol{\mu}_t) = \ldots = \tau_s^{-1} \underbrace{\left(\boldsymbol{I} - 2\boldsymbol{\Omega}_t \,\mathrm{diag}\left[\tfrac{\tau_s}{2}\boldsymbol{\lambda}^2\right]\right)}_{\circledast\circledast} \boldsymbol{\Omega}\boldsymbol{\mu}_t + \frac{\boldsymbol{C}^\top \boldsymbol{v}_t}{\widetilde{\sigma}^2_{\mathrm{obs}}}\,\delta(t - t_{\mathrm{obs}}) \;. \tag{26}$$

Again, as a sanity check, we observe the desired decay to zero because of $\circledast\circledast \to -1$, in the absence of observations.

**Continuous stream of observations.** So far, we have treated observations $\boldsymbol{v}_t$ as point observations which arrive only at distinct time points $t_{\mathrm{obs}}$ and, then, lead to "jumps" via integrating over the Dirac delta. For a complete continuous-time formulation, we choose to turn observations into a continuous input stream. When observations arrive every $\delta t$-interval and are corrupted by i.i.d. Gaussian noise of variance $\widetilde{\sigma}^2_{\mathrm{obs}}$, we can render their information content $\delta t$-independent by setting

$$\widetilde{\sigma}^2_{\mathrm{obs}} = \sigma^2_{\mathrm{obs}} / \delta t \tag{27}$$

with the alternative parameter $\sigma^2_{\mathrm{obs}}$ being independent of the observation frame rate (see, e.g., (Jazwinski, 2007; Drugowitsch et al., 2012)). Furthermore, for $\delta t \to 0$, we use that

$$\delta t \,\delta(t - t_{\mathrm{obs}}) \to 1 \tag{28}$$

because we get one Dirac delta-integration per $\delta t$ in eqn. (25) and (26) while all other variables stay (almost) constant. With these two substitutions, we obtain:

$$\partial_t \boldsymbol{\Omega}_t = \left(\tfrac{\tau_s}{2}\right)^{-1} \left(\boldsymbol{I} - \boldsymbol{\Omega}_t \,\mathrm{diag}\left[\tfrac{\tau_s}{2}\boldsymbol{\lambda}^2\right]\right) \boldsymbol{\Omega}_t + \frac{\boldsymbol{C}^\top \boldsymbol{C}}{\sigma^2_{\mathrm{obs}}} \;, \tag{29}$$

$$\partial_t (\boldsymbol{\Omega}\boldsymbol{\mu}_t) = \tau_s^{-1} (\boldsymbol{I} - 2\boldsymbol{\Omega}_t \,\mathrm{diag}\left[\tfrac{\tau_s}{2}\boldsymbol{\lambda}^2\right])\, \boldsymbol{\Omega}\boldsymbol{\mu}_t + \frac{\boldsymbol{C}^\top \boldsymbol{v}_t}{\sigma^2_{\mathrm{obs}}} \;, \tag{30}$$

with a continuous stream of observations, $\boldsymbol{v}_t$. Together, eqn. (29) and (30) solve the E-step (from **Section 3.1.1**) by re-transforming the parameters via $\boldsymbol{\Sigma}_t = \boldsymbol{\Omega}_t^{-1}$ and $\boldsymbol{\mu}_t = \boldsymbol{\Sigma}_t\,\boldsymbol{\Omega}\boldsymbol{\mu}_t$. These moments are used in the filtering posterior $p(\,\boldsymbol{s}_t \,|\, \boldsymbol{v}_{0:t}; \boldsymbol{\lambda}\,)$ of the reference online EM algorithm. The solution in terms of natural parameters is equivalent to the Kalman-Bucy filter (Kalman and Bucy, 1961) which is derived directly in terms of $\boldsymbol{\mu}_t$ and $\boldsymbol{\Sigma}_t$, as we will see in **Section 3.3.1**.

##### 3.2.3 Simultaneous online inference of motion sources and structure

We complete our derivation of online, hierarchical inference of motion sources and motion structure by casting the M-step, eqn. (20), into a recursive form and drawing the continuous-time limit. We restate eqn. (20) for reference,

$$\tfrac{\tau_s}{2}\lambda_m^2 = \frac{\frac{t\cdot\delta t}{\tau_s}\,\langle \mu_{j,m}^2 + \sigma_{j,m}^2 \rangle_j + \frac{\tau_s}{2}\nu_m \kappa_m^2}{2 + \nu_m + \frac{t\cdot\delta t}{\tau_s}} \;, \tag{31}$$

where $\mu_{j,m}$ and $\sigma^2_{j,m} = \Sigma_{j,mm}$ are the posterior parameters obtained from the E-step. Recalling that $\boldsymbol{\lambda}_t^2$ is based on the time average of $t$ discrete-time samples, we formulate $\boldsymbol{\lambda}_{t+1}^2$ as a sliding window average,

$$\boldsymbol{\lambda}_{t+1}^2 = \left(1 - \frac{1}{t}\right) \cdot \boldsymbol{\lambda}_t^2 + \frac{1}{t} \cdot \left(\frac{\tau_s}{2}\right)^{-1} \frac{\frac{t \cdot \delta t}{\tau_s} \left(\boldsymbol{\mu}_{t+1}^2 + \boldsymbol{\sigma}_{t+1}^2\right) + \frac{\tau_s}{2} \boldsymbol{\nu \kappa}^2}{2 + \boldsymbol{\nu} + \frac{t \cdot \delta t}{\tau_s}} \,, \tag{32}$$

where $(1 - \frac{1}{t})$ and $\frac{1}{t}$ are the weights for convex combination, and all vector operations are elementwise. Subtracting $\boldsymbol{\lambda}_t^2$ and dividing by inter-observation interval $\delta t$, we obtain:

$$\frac{\boldsymbol{\lambda}_{t+1}^2 - \boldsymbol{\lambda}_t^2}{\delta t} = -\frac{1}{t\,\delta t} \left( \boldsymbol{\lambda}_t^2 - \left(\frac{\tau_s}{2}\right)^{-1} \frac{\frac{t \cdot \delta t}{\tau_s} \left(\boldsymbol{\mu}_{t+1}^2 + \boldsymbol{\sigma}_{t+1}^2\right) + \frac{\tau_s}{2} \boldsymbol{\nu \kappa}^2}{2 + \boldsymbol{\nu} + \frac{t \cdot \delta t}{\tau_s}} \right) \,. \tag{33}$$

In this form, drawing the continuous-time limit is straight-forward. We let $\delta t \to 0$ while keeping $\tau_\lambda := t\,\delta t$ constant:

$$\partial_t \boldsymbol{\lambda}_t^2 = -\frac{1}{\tau_\lambda} \left( \boldsymbol{\lambda}_t^2 - \left(\frac{\tau_s}{2}\right)^{-1} \frac{\frac{\tau_\lambda}{\tau_s} \left(\boldsymbol{\mu}_t^2 + \boldsymbol{\sigma}_t^2\right) + \frac{\tau_s}{2} \boldsymbol{\nu \kappa}^2}{2 + \boldsymbol{\nu} + \frac{\tau_\lambda}{\tau_s}} \right) \,. \tag{34}$$

Time $t$ is now in continuous-time. The time constant $\tau_\lambda$ is the (continuous-time) width of the integration window and defines the minimum time scale at which significant changes of $\boldsymbol{\lambda}_t^2$ are expected to occur. From a strict algorithmic perspective of EM, we require that $\tau_\lambda \gg \tau_s$. However, we observe in computer simulations that for practical applications even small values, $\tau_\lambda \gtrapprox \tau_s$, work reliably.

**Extension to multiple spatial dimensions.** We conclude by generalizing eqn. (34) to multiple spatial dimensions via comparison of eqn. (20) and eqn. (21):

$$\partial_t \boldsymbol{\lambda}_t^2 = -\frac{1}{\tau_\lambda} \left( \boldsymbol{\lambda}_t^2 - \left(\frac{D\,\tau_s}{2}\right)^{-1} \frac{\frac{\tau_\lambda}{\tau_s} \left(\sum_{d=1}^{D} \boldsymbol{\mu}_{t,d}^2 + \boldsymbol{\sigma}_{t,d}^2\right) + \frac{\tau_s}{2} \boldsymbol{\nu \kappa}^2}{\frac{2}{D} + \boldsymbol{\nu} + \frac{\tau_\lambda}{\tau_s}} \right) \,. \tag{35}$$

Eqn. (35) is used for inferring the motion strengths $\boldsymbol{\lambda}_t^2$ in the reference online EM algorithm. The interactions of eqn. (29), (30) and (35) confirm and particularize our earlier chicken-and-egg intuition that motion sources, $\boldsymbol{s}_t$, and motion structure, $\boldsymbol{\lambda}_t$, are mutually coupled. The exact interactions are rather convoluted, and we will identify in **Section 3.3** an approximate interaction that is intuitively more accessible.

#### 3.3 Adiabatic approximation for prediction error-based inference

The inference algorithm described by eqn. (29), (30), and (35) is a nice solution from a mathematical perspective. Yet, for a theory of brain computation, it is questionable whether neural dynamics could reliably calculate 3rd-order polynomials in the variables $\boldsymbol{\Omega}\boldsymbol{\mu}_t$, $\boldsymbol{\Omega}_t$, and $\boldsymbol{\lambda}_t^2$ as demanded, for instance, by eqn. (30).

In the following, we therefore explore an alternative, approximate solution which, as we will see, considerably simplifies the involved computations while leading to almost identical results during motion structure inference. We will proceed in three steps. First, in **Section 3.3.1**, we transfer eqn. (29) + (30) back into the domain of moments, $\boldsymbol{\mu}_t$ and $\boldsymbol{\Sigma}_t$, and recover a prediction error-based update equation known as the Kalman-Bucy filter (Kalman and Bucy, 1961; Jazwinski, 2007). Then, in **Section 3.3.2**, we introduce an adiabatic approximation for the posterior covariance, $\boldsymbol{\Sigma}_t$, assuming that it has always converged to stationarity. Finally, in **Section 3.3.3**, we derive an analytical solution for the converged uncertainty for the special case of diagonal covariance. The resulting algorithm, which we term the *adiabatic observer model* in the Supporting Information and which is the online hierarchical inference model used in the main text, features interpretable dynamics on the behaviorally relevant quantities $\boldsymbol{\mu}_t$ and $\boldsymbol{\lambda}_t^2$ and relies on only quadratic computations of vector-valued variables, rather than 3rd-order computations on matrix-valued variables.

##### 3.3.1 Prediction error-based formulation

We first transform eqn. (29) into dynamics of $\boldsymbol{\Sigma}_t$:

$$\partial_t \boldsymbol{\Sigma}_t = -\boldsymbol{\Sigma}_t \left(\partial_t \boldsymbol{\Omega}_t\right) \boldsymbol{\Sigma}_t \tag{36}$$

$$= -\left(\tfrac{\tau_s}{2}\right)^{-1} \boldsymbol{\Sigma}_t + \operatorname{diag}\left[\boldsymbol{\lambda}^2\right] - \boldsymbol{\Sigma}_t \frac{\boldsymbol{C}^\top \boldsymbol{C}}{\sigma_{\text{obs}}^2} \boldsymbol{\Sigma}_t \,, \tag{37}$$

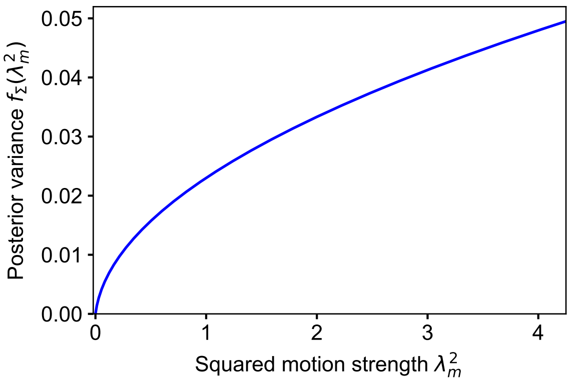

**Figure S9. Adiabatic, diagonal solution for the posterior variance.** Shown is the function $f_\Sigma(\lambda_m^2)$, given by eqn. (42). Parameters: $\tau_s = 300\,\text{ms}$, $\sigma_{\text{obs}} = 0.05$, $\|\boldsymbol{c}_m\|^2 = 4$.

and use this result for transforming eqn. (30) into dynamics of $\boldsymbol{\mu}_t$:

$$\partial_t \boldsymbol{\mu}_t = \partial_t (\boldsymbol{\Sigma}_t \boldsymbol{\Omega} \boldsymbol{\mu}_t) = (\partial_t \boldsymbol{\Sigma}_t)\, \boldsymbol{\Omega} \boldsymbol{\mu}_t + \boldsymbol{\Sigma}_t\, \partial_t \boldsymbol{\Omega} \boldsymbol{\mu}_t \tag{38}$$

$$= -\frac{\boldsymbol{\mu}_t}{\tau_s} + \boldsymbol{\Sigma}_t \boldsymbol{C}^\top \left( \frac{\boldsymbol{v}_t}{\sigma_{\text{obs}}^2} - \frac{\boldsymbol{C}\boldsymbol{\mu}_t}{\sigma_{\text{obs}}^2} \right) . \tag{39}$$

Eqn. (39) is a pretty neat equation as it reveals the “inner working” of inference as updating $\boldsymbol{\mu}_t$ with the help of prediction errors $(\boldsymbol{v}_t - \boldsymbol{C}\boldsymbol{\mu}_t)$, which are projected “up” into motion source space via $\boldsymbol{C}^\top$, and then weighted by the relative uncertainty of internal estimates vs. the uncertainty of observations, $\boldsymbol{\Sigma}_t \,/\, \sigma_{\text{obs}}^2$. In particular, in the absence of observations ($\sigma_{\text{obs}}^2 \to \infty$), the estimated mean $\boldsymbol{\mu}_t = \langle \boldsymbol{s}_t \rangle$ decays to zero with time constant $\tau_s$, as expected from the OU process. At first glance, it may seem that the inferred structure, $\boldsymbol{\lambda}^2$, plays no role in inferring $\boldsymbol{s}_t$ anymore. But actually, $\boldsymbol{\lambda}^2$ still is present in eqn. (39) indirectly through its effect on $\boldsymbol{\Sigma}_t$. We will study this indirect effect in the following.

##### 3.3.2 Convergence approximation on the posterior precision

For $\tau_\lambda > \tau_s$, we observe that $\boldsymbol{\Sigma}_t$ can be calculated directly as a function of $\boldsymbol{\lambda}_t^2$, instead of going through the hassle of integrating eqn. (37). This can be see as follows. We know that the posterior covariance $\boldsymbol{\Sigma}_t$ decays towards its stationary value with a time constant in the order of $\tau_s/2$. The stationary value itself is a dynamic equilibrium between increasing uncertainty due to diffusion (the underlying Wiener process in eqn. (2)) and decreasing uncertainty due to incoming observations (corresponds to the term $\boldsymbol{C}^\top \boldsymbol{C}/\sigma_{\text{obs}}^2$ in eqn. (29)). Notably, the stationary value does not depend on the observations, $\boldsymbol{v}_t$. This is a peculiarity of the inference task at hand which is known from Kalman filtering. This leaves $\boldsymbol{\lambda}_t^2$ as the only dynamic variable in eqn. (37) to influence the stationary point of $\boldsymbol{\Sigma}_t$. Since $\boldsymbol{\lambda}_t^2$ changes on time scales $\tau_\lambda > \tau_s$, the covariance, $\boldsymbol{\Sigma}_t$, will always have enough time to react to any change in its stationary value. This justifies treating $\boldsymbol{\Sigma}_t$ as having converged at any time, a method known in physics as adiabatic approximation.

For stationary $\boldsymbol{\Sigma}_t$, it follows from setting $\partial_t \boldsymbol{\Sigma}_t = \boldsymbol{0}$ in eqn. (37) that

$$\frac{\tau_s}{2} \boldsymbol{\Sigma} \frac{\boldsymbol{C}^\top \boldsymbol{C}}{\sigma_{\text{obs}}^2} \boldsymbol{\Sigma} + \boldsymbol{\Sigma} - \operatorname{diag}\left[\tfrac{\tau_s}{2} \boldsymbol{\lambda}^2\right] = \boldsymbol{0} . \tag{40}$$

##### 3.3.3 Analytic solution for diagonal covariance matrices

Eqn. (40) is a continuous-time algebraic Riccati equation which can, in general, be solved using eigendecompositions of an extended matrix. However, for a neural implementation, we will follow a simpler route by assuming that $\boldsymbol{\Sigma}$ is diagonal. This amounts to ignoring correlations in uncertainty about latent motion sources in the posterior distribution, for instance, during reasoning of the type: *“I have correctly decomposed all velocities in expectation, and I know my uncertainty about each motion component. But if I underestimated the flock velocity, then I likely overestimated the birds’ individual velocities.”* Only the last step of this reasoning will be ignored by dropping off-diagonal elements in $\boldsymbol{\Sigma}$. We observe in computer simulations that neglecting these posterior correlations typically has little impact on the solution.

For diagonal $\mathbf{\Sigma}$, eqn. (40) can be solved for each element $\sigma_m^2 := \Sigma_{mm}$ separately:

$$\frac{\tau_s}{2} \frac{\|\boldsymbol{c}_m\|^2}{\sigma_{\text{obs}}^2} (\sigma_m^2)^2 + \sigma_m^2 - \frac{\tau_s}{2} \lambda_m^2 = 0 \ , \tag{41}$$

where we have defined $\|\boldsymbol{c}_m\|^2 = \sum_{k=1}^{K} C_{km}^2$ to denote the vector-norm of the $m$-th column of $\boldsymbol{C}$, that is, the squared Euclidean length of the $m$-th motion component. Solving eqn. (41) is straightforward, and we denote with $f_\Sigma(\lambda_m^2)$ the resulting function for calculating $\sigma_m^2$ as a function of $\lambda_m^2$:

$$\sigma_m^2 = f_\Sigma(\lambda_m^2) = \frac{\sigma_{\text{obs}}^2}{\tau_s \|\boldsymbol{c}_m\|^2} \left( -1 + \sqrt{1 + \frac{\tau_s{}^2 \|\boldsymbol{c}_m\|^2}{\sigma_{\text{obs}}^2} \lambda_m^2} \right) \ . \tag{42}$$

$f_\Sigma$ is a monotonically increasing, non-negative function. Its graph is shown in **Fig. S9** for typical parameter values. In the limit of small motion strengths, $\lambda_m \to 0$, the variance grows quadratically in $\lambda_m$ (non-squared): $f_\Sigma(\lambda_m^2) \approx \frac{\tau_s}{2} \lambda_m^2$. For large strengths, $\lambda_m \to \infty$, the variance becomes linear in $\lambda_m$: $f_\Sigma(\lambda_m^2) \approx \frac{\sigma_{\text{obs}}}{\|\boldsymbol{c}_m\|} \lambda_m$ .

##### 3.3.4 Putting it together: neuro-friendly algorithm for online structure inference

To obtain the neuro-friendly *adiabatic observer model*, we simply plug eqn. (42) into the dynamics of $\boldsymbol{\lambda}_t^2$, given by eqn. (35), and $\boldsymbol{\mu}_t$, given by eqn. (39):

$$\partial_t \boldsymbol{\lambda}_t^2 = -\frac{1}{\tau_\lambda} \left( \boldsymbol{\lambda}_t^2 - \left( \frac{D\,\tau_s}{2} \right)^{-1} \frac{\frac{\tau_\lambda}{\tau_s} \left( \sum_{d=1}^{D} \boldsymbol{\mu}_{t,d}^2 + \boldsymbol{f}_\Sigma(\boldsymbol{\lambda}_t^2) \right) + \frac{\tau_s}{2} \boldsymbol{\nu} \boldsymbol{\kappa}^2}{\frac{2}{D} + \boldsymbol{\nu} + \frac{\tau_\lambda}{\tau_s}} \right) \ , \tag{43}$$

$$\partial_t \boldsymbol{\mu}_t = -\frac{\boldsymbol{\mu}_t}{\tau_s} + f_\Sigma(\boldsymbol{\lambda}_t^2)\, \boldsymbol{C}^{\mathsf{T}} \left( \frac{\boldsymbol{v}_t}{\sigma_{\text{obs}}^2} - \frac{\boldsymbol{C}\boldsymbol{\mu}_t}{\sigma_{\text{obs}}^2} \right) \ , \tag{44}$$

$$\text{with} \quad f_\Sigma(\lambda_m^2) = \frac{\sigma_{\text{obs}}^2}{\tau_s \|\boldsymbol{c}_m\|^2} \left( -1 + \sqrt{1 + \frac{\tau_s{}^2 \|\boldsymbol{c}_m\|^2}{\sigma_{\text{obs}}^2} \lambda_m^2} \right) \ .$$

In this vector notation, $\boldsymbol{f}_\Sigma(\boldsymbol{\lambda}_t^2)$ is evaluated elementwise, and the sum in eqn. (43) includes one evaluation of $\boldsymbol{f}_\Sigma$ for each spatial dimension $d = 1..D$. In eqn. (44), $f_\Sigma(\boldsymbol{\lambda}_t^2)$ is multiplied elementwise with the "up-projected" prediction error.

We recognize how the motion structure, $\boldsymbol{\lambda}_t$, controls the gating function, $f_\Sigma(\boldsymbol{\lambda}_t^2)$, for performing the credit assignment of the prediction errors, in eqn. (44). Furthermore, we note that also the posterior uncertainty can be recovered at any time since $\sigma_{t,m}^2 = f_\Sigma(\lambda_{t,m}^2)$.

For $D = 1$, eqn. (43) is eqn. (1) from the main text, eqn. (44) is eqn. (2), and eqn. (42) is eqn. (3).

##### 3.3.5 A pleasant note on inference of rotational and radial motion

The approximations introduced for the adiabatic observer model hold, remarkably and importantly, also for the biologically relevant case of radial and rotational motion (cf. **Section 2.5** and **Fig. S8** for the generative model).

To illustrate this, consider the case of two motion sources, radial $s_{\text{rad}}(t)$ and rotational $s_{\text{rot}}(t)$, with motion features given by eqn. (6). The motion features $\boldsymbol{C}$ depend on the receptive field locations with parameters $R_k$ and $\vartheta_k$. For inferring the radial component, $s_{\text{rad}}(t)$, all receptive fields are equally informative, irrespective of their eccentricity, $R_k$. Accordingly, eqn. (44) weighs all radial prediction errors equally, as expressed by the $R_k$-independence of the radial row in $\boldsymbol{C}^{\mathsf{T}}$. This is different when estimating rotational motion, $s_{\text{rot}}(t)$. Here, prediction errors measured near the fovea (small $R_k$) contribute only little to $\partial_t \mu_{\text{rot}}$: for small $R_k$, we expect to observe a small rotational motion energy via $\boldsymbol{C}\boldsymbol{\mu}_t$, such that the "Cartesian" observation noise of size $\sigma_{\text{obs}}$ makes the stimulus virtually uninformative about the rotational velocity $\dot{\varphi}$. The scaling of $\boldsymbol{C}^{\mathsf{T}}$ with $R_k$ accounts for that. Receptive fields far away from the fovea, in contrast, predict a strong rotational velocity via $\boldsymbol{C}\boldsymbol{\mu}_t$, such that noise of size $\sigma_{\text{obs}}$ (in Cartesian space) has only a minor impact on estimating $\mu_{\text{rot}}$. Accordingly, the scaling of $\boldsymbol{C}^{\mathsf{T}}$ with $R_k$ assigns a higher weight to peripheral receptive fields for estimating rotational motion.

As a final remark, the above example assumed the observation noise $\sigma_{\text{obs}}$ to be constant across all receptive fields. Eqn. (44) naturally supports extensions to heterogeneous observation noise because all local prediction errors are measured in units of their local noise.

### 4. Extensions of the online model

#### 4.1 Non-isotropic observation noise and missing observations

For the main manuscript, we have assumed the observation noise, $\sigma_{\text{obs}}$, to be a constant across time, $t$, observed features, $k$, and spatial dimensions, $d$. In real-world scenes, the observation noise could change along all those indices. An object could be occluded, or otherwise temporarily invisible, leading to $1/\sigma_{\text{obs}}^2 = 0$. Different objects might have different observation noise, e.g., due to different visual contrast. The aperture problem could render local velocity signals ambiguous: e.g., what is the direction of motion for a straight line that is larger than the aperture? This could be modeled by small observation noise perpendicular to the line, and large noise parallel to the line. The so-constructed diagonal covariance matrix, $\boldsymbol{\Sigma}_{\text{diag}}$, is then rotated as per $\boldsymbol{\Sigma}_{\text{xy}} = \boldsymbol{Q}^\top \boldsymbol{\Sigma}_{\text{diag}} \boldsymbol{Q}$ with rotation matrix, $\boldsymbol{Q}$, into the canonical $x$-$y$-coordinate frame.

The above extensions of the observation noise are supported by our online model, as long as changes in $\sigma_{\text{obs}}^2$ occur slower than $\tau_s$ (so the adiabatic approximation remains valid). In eqn. (44), $\frac{1}{\sigma_{\text{obs}}^2}$ is extended to have different elements, $\frac{1}{\sigma_{\text{obs},k,d}^2}$, and is multiplied elementwise with $v_{t,k,d}$ and $(\boldsymbol{C}\boldsymbol{\mu})_{t,k,d}$. When calculating $f_\Sigma$, replace

$$\frac{\|\boldsymbol{c}_m\|^2}{\sigma_{\text{obs}}^2} \qquad \text{by} \qquad \left( \boldsymbol{C}^\top \operatorname{diag}\left[ \frac{\mathbf{1}}{\boldsymbol{\sigma}_{\text{obs}}^2} \right] \boldsymbol{C} \right)_{mm} \tag{45}$$

for each spatial dimension. In eqn. (43), the summation then runs over the different spatial dimensions of $f_\Sigma$. The idea that $\frac{1}{\sigma_{\text{obs}}^2}$ is vector-valued is also used in the network implementation in **Section 5**.

A note on the provided Python code package: In the code for the network, $\boldsymbol{\sigma}_{\text{obs}}^2$ can currently have different values for every input velocity $v_k$, but is assumed to be (a) identical in both spatial dimensions, and (b) not to change over time. The code for the algorithm is less restrictive by supporting temporary masking of inputs (presented via `class ObservationGeneratorVelo`; leading to posterior variance $\sigma_m^2 = f_\Sigma(\lambda_m^2) = \frac{\tau_s}{2}\lambda_m^2$ if all dependent objects are invisible according to eqn. (41)) and different noise in spatial dimensions.

#### 4.2 Heterogeneous time constants

Different motion components, $s_m$, might have different time-constants, $\tau_s$, for typical changes in speed and direction to occur. An extension to a vector $\boldsymbol{\tau}_s = (\tau_{s,1}, .., \tau_{s,M})$ is straightforward by replacing all occurences of $\tau_s$ and $1/\tau_s$ by $\operatorname{diag}[\boldsymbol{\tau_s}]$ and $\operatorname{diag}[\boldsymbol{\tau_s}^{-1}]$, respectively. The code package supports heterogeneous time constants for both the algorithm and the network.

#### 4.3 Interaction priors capturing feature compatibility

Some motion components may be unlikely to occur together. Consider, for example, two cluster components in $\boldsymbol{C}$ which are overlapping but do not contain one another: the simultaneous occurrence of the two clusters would not be compatible with a tree structure. We can accommodate such incompatibility with the help of an *interaction prior*.

In the following, we outline how interaction priors can be included in the theory, and how they will affect the inference process. We endow the $\boldsymbol{\lambda}^2$-prior from eqn. (16) with an interaction term:

$$p(\boldsymbol{\lambda}^2; \boldsymbol{\nu}, \boldsymbol{\kappa}^2) \propto \left[ \prod_{m=1}^{M} \mathcal{I}\chi(\lambda_m^2; \nu_m, \kappa_m^2) \right] \cdot e^{-\frac{1}{2}(\boldsymbol{\lambda}^2)^\top \boldsymbol{J} (\boldsymbol{\lambda}^2)}, \tag{46}$$

where the interaction matrix $\boldsymbol{J} \in \mathbb{R}^{M \times M}$ is a symmetric, zero-diagonal matrix that models feature incompatibility. For instance, positive values, $J_{ml} = J_{lm} > 0$, describe a (soft) incompatibility between the $m^{\text{th}}$ and $l^{\text{th}}$ motion component.

In the derivation of the M-step, the interaction prior leads to an additional term $(-\boldsymbol{J}\boldsymbol{\lambda})_m$ on the right-hand side of eqn. (17).[4] For the optimum in eqn. (19), this leads to the following equation (using vector notation and covering multiple spatial dimensions):

$$D\,\frac{\tau_s}{2} \left( \boldsymbol{I} + \frac{2}{2/D + \boldsymbol{\nu} + t} \operatorname{diag}\left[(\boldsymbol{\lambda}^2)^2\right] \boldsymbol{J} \right) \boldsymbol{\lambda}^2 = \frac{t \sum_d \langle \boldsymbol{\mu}_j^2 + \sigma_j^2 \rangle_j + \frac{\tau_s}{2} \boldsymbol{\nu} \boldsymbol{\kappa}^2}{2/D + \boldsymbol{\nu} + t}\ . \tag{47}$$

[4]Note that the normalization of the distribution in eqn. (46) will not depend on $\lambda^2$, and thus will play no role in the M-step, which uses $\lambda^2$-derivatives of $\log p(\lambda^2)$.

For small values of $\|\boldsymbol{J}\|$ or, similarly, large values of $t$ ($=\tau_\lambda/\tau_s$ in continuous-time), the matrix $\widetilde{\boldsymbol{J}} := \left(\boldsymbol{I} + \frac{2}{2/D+\nu+t}\mathrm{diag}\left[(\boldsymbol{\lambda}^2)^2\right]\boldsymbol{J}\right)$ is invertible with the approximate inverse $\widetilde{\boldsymbol{J}}^{-1} \approx \left(\boldsymbol{I} - \frac{2}{2/D+\nu+t}\mathrm{diag}\left[(\boldsymbol{\lambda}^2)^2\right]\boldsymbol{J}\right)$. We can therefore move this matrix to the right-hand side and follow the derivation for online inference without interaction priors. This leads to the following equivalent of eqn. (43):

$$\partial_t \boldsymbol{\lambda}_t^2 = -\frac{1}{\tau_\lambda}\left(\boldsymbol{\lambda}_t^2 - \left(\frac{D\,\tau_s}{2}\right)^{-1}\left(\boldsymbol{I} - \frac{2}{\frac{2}{D}+\boldsymbol{\nu}+\frac{\tau_\lambda}{\tau_s}}\mathrm{diag}\left[(\boldsymbol{\lambda}^2)^2\right]\boldsymbol{J}\right)\frac{\frac{\tau_\lambda}{\tau_s}\left(\sum_{d=1}^{D}\boldsymbol{\mu}_{t,d}^2 + \boldsymbol{f}_\Sigma(\boldsymbol{\lambda}_t^2)\right) + \frac{\tau_s}{2}\boldsymbol{\nu}\boldsymbol{\kappa}^2}{\frac{2}{D}+\boldsymbol{\nu}+\frac{\tau_\lambda}{\tau_s}}\right) \,. \tag{48}$$

The only difference to eqn. (43) is that the target values on the right hand side are mixed together via $\widetilde{\boldsymbol{J}}^{-1}$. This gives rise to quite intuitive dynamics: if two motion components are incompatible, they mutually subtract their respective (independent) target values from another, thereby slightly changing the motion structure in which the E-step will interpret future input and, ultimately, leading to soft winner-takes-all competition. The term $(\mathrm{diag}\left[(\boldsymbol{\lambda}^2)^2\right])$ limits the competition to those components that are significantly different from zero.

#### 4.4 Detecting motion components that had decayed to baseline

If motion components are not detected in the structure for a longer time, the associated strength, $\lambda_m$, will decay to zero. Since $f_\Sigma(0)=0$, this will prohibit future detection of said motion component. This issue can be addressed in two ways. In a biological agent, noise in the nervous system will lead to small fluctuations in the encoded value of $\boldsymbol{\lambda}$, thereby "probing" the presence of motion components simply via noisy deviations from $\lambda_m=0$. Alternatively, a more principled solution exploits the hyperparameters, $\nu_m$ and $\kappa_m$, in eqn. (16) to prevent $\lambda_m$ from decaying to zero, e.g., by choosing $\nu_m=1$ and $\kappa_m=0.1$. This follows the intuition that pseudo-observations in support of $\lambda_m=\kappa_m$ had been observed for a duration $\nu_m\,\tau_s$.

#### 4.5 Learning the motion components on long time-scales

While not being the focus of this work, we briefly touch upon the question of how the motion components, $\boldsymbol{C}$, could be learned online from observations in an unsupervised manner. To this end, we follow a similar EM scheme as for inferring $\boldsymbol{\lambda}$ and note that in eqn. (13) only the quadratic term depends on $\boldsymbol{C}$. For the M-step, however, instead of maximizing $Q$ directly, we perform gradient ascent with respect to $\boldsymbol{C}$:

$$\nabla_{\boldsymbol{C}} Q_t(\boldsymbol{C}) = \langle \nabla_{\boldsymbol{C}}[-\frac{1}{2\sigma_{\mathrm{obs}}^2}(\boldsymbol{v}_t - \boldsymbol{C}\boldsymbol{s}_t)^\top(\boldsymbol{v}_t - \boldsymbol{C}\boldsymbol{s}_t)]\rangle_{p(\boldsymbol{s}_t\,|\,\boldsymbol{v}_{0:t})} \tag{49}$$

$$= \frac{1}{\sigma_{\mathrm{obs}}^2}\langle \boldsymbol{v}_t\boldsymbol{s}_t^\top - \boldsymbol{C}\,(\boldsymbol{s}_t\boldsymbol{s}_t^\top)\rangle_{p(\boldsymbol{s}_t\,|\,\boldsymbol{v}_{0:t})} \tag{50}$$

$$= \frac{1}{\sigma_{\mathrm{obs}}^2}\left(\boldsymbol{v}_t\boldsymbol{\mu}_t^\top - \boldsymbol{C}\left(\boldsymbol{\mu}_t\boldsymbol{\mu}_t^\top + \boldsymbol{\Sigma}_t\right)\right) \,. \tag{51}$$

This gradient establishes the intuition to compare the observed covariance between inputs and motion components against their expected covariance.

Furthermore, we note that the parameterization of motion structure via $\boldsymbol{C}$ and $\boldsymbol{\lambda}$ leaves an invariance: any scaling of $\boldsymbol{\lambda}$ can be compensated by an inverse scaling of $\boldsymbol{C}$. Due to the sparsity prior on $\boldsymbol{\lambda}$, which favors small values, this bears the risk of unbounded growth in $\boldsymbol{C}$. We can address this risk by imposing a regularizing prior on $\boldsymbol{C}$, e.g., a Laplace prior or a Gaussian prior, such that the system finds a balance between small $\boldsymbol{\lambda}$ and small $\|\boldsymbol{C}\|$. We incorporate the regularizer in the gradient-based update rule with the aim to balance prior and likelihood such that the likelihood is weighted to contribute $N_C$ independent samples:

$$\partial_t \boldsymbol{C} = \eta_C\left[\frac{1}{\sigma_{\mathrm{obs}}^2}\left(\boldsymbol{v}_t\boldsymbol{\mu}_t^\top - \boldsymbol{C}\left(\boldsymbol{\mu}_t\boldsymbol{\mu}_t^\top + \boldsymbol{\Sigma}_t\right)\right) + \frac{1}{N_C}\nabla_{\boldsymbol{C}}\log p(\boldsymbol{C})\right] \,. \tag{52}$$

Examples for the regularizer are $\nabla_{\boldsymbol{C}}\log p(\boldsymbol{C}) = -\mathrm{sign}(\boldsymbol{C})/b$ for a Laplace prior and $\nabla_{\boldsymbol{C}}\log p(\boldsymbol{C}) = -\boldsymbol{C}/b$ for a Gaussian prior. For the online M-step, both $\boldsymbol{\mu}_t$ and $\boldsymbol{\lambda}_t$ must have time to adapt to any changes in $\boldsymbol{C}$. This is ensured by requiring that the learning rate, $\eta_C$, is small enough to average over a large number of independent samples, i.e., $\eta_C \ll 1/\tau_\lambda$. The small learning rate emphasizes how we think of $\boldsymbol{C}$-optimization as a long-term learning process of a "feature dictionary".

Eqn. (52) offers a path to learning $\boldsymbol{C}$ via an online EM algorithm. In a network implementation with linear population codes, however, it remains unclear how the update could be communicated to synapses: $\boldsymbol{C}$ determines many synaptic weights in the network. Finding encoding and decoding vectors of the variables to support simple plasticity rules is future work.

### 5. Neural network implementation

*How could biological recurrent neural networks implement online motion structure inference?* In light of the theory developed in **Section 3.3**, we will operationalize this question by implementing eqn. (43) and eqn. (44).

While we will strive to incorporate salient properties from motion sensitive brain areas, the exact computational mechanisms underlying many experimental findings are still elusive. Thus, inevitably, several modeling assumptions have to be made. These assumptions are presented in **Section 5.1**. We then discuss, in **Section 5.2**, which variables—input and latent—we choose to be linearly decodable by downstream populations, and, in **Section 5.3**, express the adiabatic observer model in terms of these variables. For performing the required computations on these variables, we will extend the ideas developed in (Beck, Latham, and Pouget, 2011), in **Section 5.4**, to a systematic theory of neural integration of high-dimensional linear and quadratic differential equations. In **Section 5.5**, we will apply the theory to derive a rate-based recurrent neural network model for online motion structure inference. Finally, in **Section 5.6**, we introduce—as an example for the computer simulations—a neural encoding model of input variables that captures many properties of middle temporal visual area (MT) while staying mathematically tractable with regard to its computational function.

#### 5.1 Aims and assumptions

We view our network model as a starting point for an experiment–theory loop. Some neural response properties will be rather general and could be tested in experiments directly. Others will be more specific and could guide targeted experiments. In any case, we expect that many aspects of this initial model will be revised and refined in the process.

For the model, we make three assumptions:

- **Rate-based network.** We assume that all information is conveyed in the neuronal firing rates. Thus, no exact spike-timing is considered. Further, we will allow negative firing rates—think of them as negative deviations from a baseline value.
- **Linear and quadratic operations.** We assume that neurons can integrate their synaptic inputs in two ways: linearly and quadratically. Specifically, we assume that the dynamics of the firing rate of a neuron (or small population) $i$ takes the form,
$$\tau_i \partial_t r_i = -r_i + f_i(\boldsymbol{w}_i^\top \boldsymbol{r} + \boldsymbol{r}^\top \boldsymbol{Q}^{(i)} \boldsymbol{r} + b_i) \ , \tag{53}$$
with time constant $\tau_i$, a potentially non-linear activation function $f_i$, weight vector $\boldsymbol{w}_i^\top$, quadratic interaction matrix $\boldsymbol{Q}^{(i)}$, and bias $b_i$. In the main text, we had omitted the (per-neuron) bias, $b_i$, because it can be absorbed in $f_i$. For the following formal derivation, we make the bias explicit for clarity, and, as we will see, this leads to activation functions, $f_i$, which are different only on a per-population basis. Eqn. (53) is a standard form for rate-based network models (Dayan and Abbott, 2001), and quadratic interactions are commonly used in neural network modeling (Salinas and Abbott, 1996; Beck, Latham, and Pouget, 2011).
- **Linear decoding of task-relevant variables.** A subset of variables, especially those which are relevant for actions and decision making, can be read out linearly by downstream populations. We will specify the subset of variables in the next section.

#### 5.2 Linearly decodable variables

Inspecting eqn. (43) and eqn. (44), an elegant decomposition into basic operations (that is, addition, linear and quadratic multiplication) employs the following variables:

$$\boldsymbol{\mu}_t \ , \qquad \boldsymbol{\lambda}_t^2 \ , \qquad \frac{\boldsymbol{v}_t}{\sigma_{\text{obs}}^2} \ , \qquad \frac{1}{\sigma_{\text{obs}}^2} \ , \qquad \underbrace{\left( \frac{\boldsymbol{v}_t}{\sigma_{\text{obs}}^2} - \frac{\boldsymbol{C}\boldsymbol{\mu}_t}{\sigma_{\text{obs}}^2} \right)}_{\text{Pred. err. } \boldsymbol{\epsilon}_t} , \qquad f_\Sigma(\boldsymbol{\lambda}_t^2) \ . \tag{54}$$

The first three variables, $\boldsymbol{\mu}_t$, $\boldsymbol{\lambda}_t^2$, $\boldsymbol{v}_t$, are directly related to the task of decomposing visual scenes. Further, we include the observation noise, $1/\sigma_{\text{obs}}^2$, as an input variable (rather than treating it as a constant) to accommodate the extended theory presented in **Section 4.1**, permitting the network to handle, for instance, transient occlusion of objects. Of course, the value of $\sigma_{\text{obs}}^2$ could also be a constant in the network. The prediction error, $\boldsymbol{\epsilon}_t$, is an auxiliary variable to restrict the complexity of operations to being at most quadratic. (We will see in **Section 5.4** that the computational complexity is directly inherited by the neural dynamics.) Finally, the posterior variance, $f_\Sigma(\boldsymbol{\lambda}_t^2)$, which is a hallmark of Bayesian computation, is required for motion structure decomposition and, potentially, for Bayesian decision making.

The variables listed in eqn. (54) are assumed to be linearly decodable, that is, they can be read out from neural activity (at the example of $\boldsymbol{\mu}_t$) via

$$\boldsymbol{\mu}_t = \boldsymbol{A}^\mu \boldsymbol{r}_t \ , \tag{55}$$

with some readout matrix $\boldsymbol{A}^\mu$. The other variables maintain corresponding matrices $\boldsymbol{A}^\lambda$, $\boldsymbol{A}^v$, $\boldsymbol{A}^\sigma$, $\boldsymbol{A}^\epsilon$, and $\boldsymbol{A}^\Sigma$, respectively. Here, $\boldsymbol{r}_t$ are the instantaneous firing rates of a population of neurons that encode $\boldsymbol{\mu}_t$. Multiple variables can be encoded by the same neural population.

#### 5.3 Motion structure inference via at-most quadratic operations

First, we observe that eqn. (43) and eqn. (44) almost exclusively contain linear and quadratic terms when expressed in the variables of eqn. (54):

$$\partial_t \boldsymbol{\lambda}_t^2 = -\frac{1}{\tau_\lambda}\boldsymbol{\lambda}_t^2 + \frac{2}{D\,\tau_s\,\tau_\lambda\left(\frac{2}{D} + \boldsymbol{\nu} + \frac{\tau_\lambda}{\tau_s}\right)}\left(\frac{\tau_\lambda}{\tau_s}\sum_{d=1}^{D}\boldsymbol{\mu}_{t,d}\odot\boldsymbol{\mu}_{t,d} + \frac{D\,\tau_\lambda}{\tau_s}\boldsymbol{f}_\Sigma(\boldsymbol{\lambda}_t^2) + \tfrac{\tau_s}{2}\boldsymbol{\nu}\boldsymbol{\kappa}^2\right) \ , \tag{56}$$

$$\partial_t \boldsymbol{\mu}_t = -\frac{1}{\tau_s}\boldsymbol{\mu}_t + \boldsymbol{f}_\Sigma(\boldsymbol{\lambda}_t^2)\odot \boldsymbol{C}^\top\boldsymbol{\epsilon}_t \ , \tag{57}$$

$$\partial_t \boldsymbol{\epsilon}_t = -\frac{1}{\tau_\epsilon}\left(\boldsymbol{\epsilon}_t - \frac{\boldsymbol{v}_t}{\boldsymbol{\sigma}_{\text{obs}}^2} + \frac{1}{\boldsymbol{\sigma}_{\text{obs}}^2}\odot\boldsymbol{C}\boldsymbol{\mu}_t\right) = -\frac{1}{\tau_\epsilon}\boldsymbol{\epsilon}_t + \frac{1}{\tau_\epsilon}\frac{\boldsymbol{v}_t}{\boldsymbol{\sigma}_{\text{obs}}^2} - \frac{1}{\tau_\epsilon}\frac{1}{\boldsymbol{\sigma}_{\text{obs}}^2}\odot\boldsymbol{C}\boldsymbol{\mu}_t \ , \tag{58}$$

where we have made elementwise multiplication explicit via the $\odot$-operator, and moved the prediction error, $\boldsymbol{\epsilon}_t$, into a separate dynamic equation with time constant $\tau_\epsilon$. For this separation to maintain faithful results, we require that $\tau_\epsilon < \tau_s$, such that the prediction error can react to changes in $\boldsymbol{\mu}_t$. The only variable in eqn. (54) that cannot be calculated within this scheme is $f_\Sigma(\boldsymbol{\lambda}^2)$, which contains a square root, and, thus, has to be addressed separately, below. The variables $\boldsymbol{v}_t/\sigma_{\text{obs}}^2$ and $1/\sigma_{\text{obs}}^2$ are the input variables that are fed into the system.

#### 5.4 Neural dynamics for integrating linear and quadratic differential equations

We will now establish how linear and quadratic dynamics of latent variables, such as eqn. (56) – (58), can be integrated in neural space. What follows is basically a clearly structured generalization of (Beck, Latham, and Pouget, 2011).

**Notation in the algorithmic domain.** Inevitably, some notation has to be introduced for addressing all of the above dynamics in both the *algorithmic domain* (i.e., dynamics of variables) and the *network domain* (i.e., dynamics of neuronal firing rates). To keep the presentation general, we will adopt variable-dynamics of the generic form

$$\partial_t \boldsymbol{z} = \boldsymbol{y}\underline{\boldsymbol{Q}}\boldsymbol{x} + \boldsymbol{W}\boldsymbol{x} + \boldsymbol{b} \qquad \text{with} \qquad (\boldsymbol{y}\underline{\boldsymbol{Q}}\boldsymbol{x})_i \stackrel{\text{def}}{=} \sum_{j,k} \underline{Q}_{ijk}\, y_j\, x_k \ . \tag{59}$$

Here, $\boldsymbol{z}$, $\boldsymbol{y}$, and $\boldsymbol{x}$ denote vector-valued variables. Further, $\boldsymbol{b}$ is a vector-valued additive constant, $\boldsymbol{W}$ a matrix, and $\underline{\boldsymbol{Q}}$ a 3rd-order tensor, for which we establish the notation of small, capital, and underlined capital letters, respectively.

Note that expressions with elementwise multiplication are covered by the tensors. For instance, using Einstein summation convention,

$$(\boldsymbol{y}\odot\boldsymbol{x})_i = y_i\,x_i = \delta_{ij}\delta_{ik}y_j\,x_k = (\boldsymbol{y}\underline{\boldsymbol{Q}}\boldsymbol{x})_i \qquad \text{with} \qquad \underline{Q}_{ijk} = \delta_{ij}\delta_{ik} \ , \tag{60}$$

$$\left[\boldsymbol{W}^1(\boldsymbol{y}\odot\boldsymbol{W}^2\boldsymbol{x})\right]_i = W^1_{ij}(\boldsymbol{y}\odot\boldsymbol{W}^2\boldsymbol{x})_j = W^1_{ij}(y_j W^2_{jk}x_k) = (\boldsymbol{y}\underline{\boldsymbol{Q}}\boldsymbol{x})_i \qquad \text{with} \qquad \underline{Q}_{ijk} = W^1_{ij}W^2_{jk} \ , \tag{61}$$

$$\left[(\boldsymbol{W}^1\boldsymbol{y})\odot(\boldsymbol{W}^2\boldsymbol{x})\right]_i = W^1_{ij}y_j W^2_{ik}x_k = (\boldsymbol{y}\underline{\boldsymbol{Q}}\boldsymbol{x})_i \qquad \text{with} \qquad \underline{Q}_{ijk} = W^1_{ij}W^2_{ik} \ . \tag{62}$$

Thus, all algorithmic dynamics in eqn. (56) – (58) are of form eqn. (59).

**Notation in the network domain.** We now turn to the question of how neuronal populations can calculate dynamics of the form in eqn. (59) when the involved variables are linearly decodable, that is, when $\boldsymbol{z} = \boldsymbol{A}^z \boldsymbol{r}^z$. Refining the notation in eqn. (55), we will make explicit which population $\boldsymbol{r}^z$ encodes variable $\boldsymbol{z}$ and suppress the time dependence in $\boldsymbol{r}_t^z$. Again, we emphasize that differently denoted populations, e.g., $\boldsymbol{r}^z$ and $\boldsymbol{r}^x$, can and often will refer to the same population—the refined notation simply gives us the flexibility to cover various cases.

Following (Beck, Latham, and Pouget, 2011), we will further make use of what is called the *adjoint matrix* $\boldsymbol{A}^{z\dagger}$ of matrix $\boldsymbol{A}^z$, which is characterized by $\boldsymbol{A}^z \boldsymbol{A}^{z\dagger} = \boldsymbol{I}$. Such right-inverse, albeit not unique, always exists if the rows of $\boldsymbol{A}^z$ are linearly independent, which is commonly fulfilled when the number of neurons exceeds the number of variables. If variables $\boldsymbol{z}$ and $\boldsymbol{x}$ are encoded by the same populations of neurons, we further require that $\boldsymbol{A}^z \boldsymbol{A}^{x\dagger} = \boldsymbol{A}^x \boldsymbol{A}^{z\dagger} = \boldsymbol{0}$. As long as these orthogonality conditions are satisfied, the exact form of the matrices $\boldsymbol{A}$ is arbitrary, from a mathematical point of view.

As we will show in the next paragraph, quadratic, linear, and constant terms in the algorithmic domain, eqn. (59), translate one-to-one into quadratic, linear and constant terms in the network domain. We therefore establish the notation $\overline{\boldsymbol{Q}}$, $\overline{\boldsymbol{W}}$, and $\overline{\boldsymbol{b}}$ (with an overbar) to refer to 3rd-order tensors, matrices and biases in the neural domain, respectively. These are exactly the function arguments that we had deemed feasible in eqn. (53) (there introduced without the overbar).

**Neural dynamics.** The neural dynamics for implementing each of the computations in eqn. (59) are as follows.

Quadratic terms: $\partial_t \boldsymbol{z} = \boldsymbol{y}\boldsymbol{Q}\boldsymbol{x}$ is implemented via

$$\partial_t \boldsymbol{r}^z = \boldsymbol{r}^y \overline{\boldsymbol{Q}} \boldsymbol{r}^x \qquad \text{with} \qquad \overline{Q}_{ijk} \stackrel{\text{def}}{=} A^{z\dagger}_{i\alpha} Q_{\alpha\beta\gamma} A^y_{\beta j} A^x_{\gamma k} \ , \qquad \text{in short:} \qquad \overline{\boldsymbol{Q}} \stackrel{\text{def}}{=} \boldsymbol{A}^{z\dagger} \left( \boldsymbol{Q} \boldsymbol{A}^y \boldsymbol{A}^x \right) \ . \tag{63}$$

Proof:

$$(\partial_t \boldsymbol{z})_i = (\partial_t \boldsymbol{A}^z \boldsymbol{r}^z)_i = (\boldsymbol{A}^z \partial_t \boldsymbol{r}^z)_i = A^z_{ij} (\boldsymbol{r}^y \overline{\boldsymbol{Q}} \boldsymbol{r}^x)_j = A^z_{ij} \overline{Q}_{jkl} r^y_k r^x_l = A^z_{ij} A^{z\dagger}_{j\alpha} Q_{\alpha\beta\gamma} A^y_{\beta k} A^x_{\gamma l} r^y_k r^x_l = \delta_{i\alpha} Q_{\alpha\beta\gamma} y_\beta x_\gamma = (\boldsymbol{y}\boldsymbol{Q}\boldsymbol{x})_i \ . \tag{64}$$

Linear terms: $\partial_t \boldsymbol{z} = \boldsymbol{W}\boldsymbol{x}$ is implemented via

$$\partial_t \boldsymbol{r}^z = \overline{\boldsymbol{W}} \boldsymbol{r}^x \qquad \text{with} \qquad \overline{\boldsymbol{W}} = \boldsymbol{A}^{z\dagger} \boldsymbol{W} \boldsymbol{A}^x \ . \tag{65}$$

Proof:

$$\partial_t \boldsymbol{z} = \boldsymbol{A}^z \partial_t \boldsymbol{r}^z = \boldsymbol{A}^z \overline{\boldsymbol{W}} \boldsymbol{r}^x = \boldsymbol{A}^z \boldsymbol{A}^{z\dagger} \boldsymbol{W} \boldsymbol{A}^x \boldsymbol{r}^x = \boldsymbol{W}\boldsymbol{x} \ . \tag{66}$$

Constant terms: $\partial_t \boldsymbol{z} = \boldsymbol{b}$ is implemented via

$$\partial_t \boldsymbol{r}^z = \overline{\boldsymbol{b}} \qquad \text{with} \qquad \overline{\boldsymbol{b}} = \boldsymbol{A}^{z\dagger} \boldsymbol{b} \ . \tag{67}$$

Proof:

$$\partial_t \boldsymbol{z} = \boldsymbol{A}^z \partial_t \boldsymbol{r}^z = \boldsymbol{A}^z \overline{\boldsymbol{b}} = \boldsymbol{A}^z \boldsymbol{A}^{z\dagger} \boldsymbol{b} = \boldsymbol{b} \ . \tag{68}$$

Linearity: Neural dynamics for linear combinations, e.g., $\partial_t \boldsymbol{z} = \boldsymbol{W}^1 \boldsymbol{x} + \boldsymbol{W}^2 \boldsymbol{y}$ are simply the sum of the individual terms, e.g., $\partial_t \boldsymbol{r}^z = \overline{\boldsymbol{W}^1} \boldsymbol{r}^x + \overline{\boldsymbol{W}^2} \boldsymbol{r}^y$. The proof follows directly from the linearity of $\boldsymbol{A}^z$.

Shared populations: Neural dynamics of variables encoded by the same population, e.g., $\boldsymbol{r} = \boldsymbol{r}^z = \boldsymbol{r}^x$, do not interfere. The proof follows from the orthogonality $\boldsymbol{A}^z \boldsymbol{A}^{x\dagger} = \boldsymbol{A}^x \boldsymbol{A}^{z\dagger} = \boldsymbol{0}$ and from observing that every term in the above neural dynamics is led by an adjoint matrix $\boldsymbol{A}^{z\dagger}$ or $\boldsymbol{A}^{x\dagger}$. Therefore, neural dynamics inducing changes in $\boldsymbol{z}$ do not convey any changes in $\boldsymbol{x}$, and vice versa.

#### 5.5 Recurrent network model for online motion structure inference

**Section 5.4** provides us with a straight-forward recipe for implementing eqn. (56) – (58) in a neural network. To keep the network as general as possible, the input variables, $1/\sigma^2_{\text{obs}}$ and $\boldsymbol{v}/\sigma^2_{\text{obs}}$, are encoded by an *input population*, $\boldsymbol{r}_{\text{inp}}$. The latent variables, $\boldsymbol{\lambda}^2$, $\boldsymbol{\mu}$ and $\boldsymbol{\epsilon}$, are encoded by a *distributed population*, $\boldsymbol{r}_{\text{dis}}$. Both the input and distributed population employ a distributed code with arbitrary readout matrices $\boldsymbol{A}$ obeying the orthonormality conditions stated in **Section 5.4**. For the distributed population, the activation function, $f_i$, in eqn. (53) is simply the identity

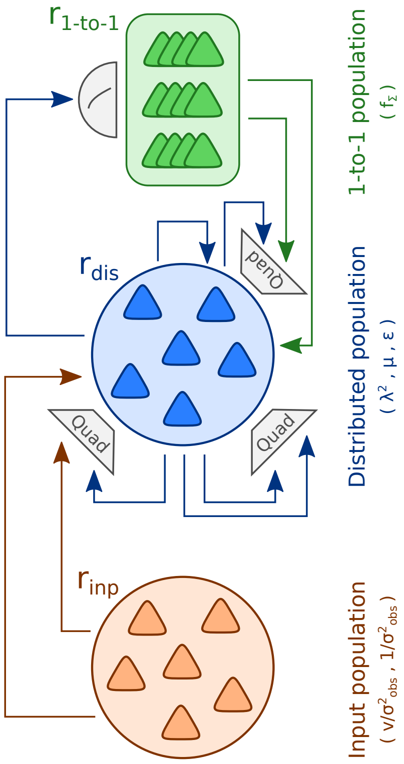

**Figure S10. Network model for motion structure inference.** The network is composed of three neuronal populations. The input population, $\boldsymbol{r}_{\text{inp}}$, encodes the input variables, $1/\sigma^2_{\text{obs}}$ and $v/\sigma^2_{\text{obs}}$, as a distributed code. The distributed population, $\boldsymbol{r}_{\text{dis}}$, encodes the latent variables, $\boldsymbol{\lambda}^2$, $\boldsymbol{\mu}$ and $\boldsymbol{\epsilon}$, as a distributed code. The one-to-one population, $\boldsymbol{r}_{\text{1-to-1}}$, encodes the latent posterior uncertainty, $f_\Sigma(\boldsymbol{\lambda}^2)$, as a one-to-one code. All of these variables can be read out linearly from the network firing rate, at any time. Synaptic connections within and between populations mediate linear (indicated as arrows) and quadratic (indicated as "Quad" boxes) interactions. The non-linear function $f_\Sigma$ is implemented by a leaky integrate-and-fire type response (indicated by the half-circle).
Overall, the network implements eqn. (56) – (58) of the algorithmic domain and, thereby, emulates the adiabatic observer model given by eqn. (43) + (44).

function. We refrain from restating the exact neural dynamics here because they are obtained directly by translating the terms in eqn. (56) – (58) into their neural counterparts by means of eqn. (63), (65), and (67).

For a functioning network model, however, two pieces are missing: the input code, and handling of the function $f_\Sigma(\boldsymbol{\lambda}_t^2)$ as was promised in **Section 5.3**. These two pieces are discussed next.

**Connecting the input.** The input is fed into the network externally and is thus by definition not controlled by internal dynamics of the network. Nonetheless, the activity $\boldsymbol{r}_{\text{inp}}(\boldsymbol{v}/\sigma^2_{\text{obs}}, 1/\sigma^2_{\text{obs}})$, which is a function of the input variables, is required to support linearly decoding $\boldsymbol{v}/\sigma^2_{\text{obs}}$ and $1/\sigma^2_{\text{obs}}$ via known readout matrices $\boldsymbol{A}^v$ and $\boldsymbol{A}^\sigma$. Note that no adjoint matrices are required for the input. While any valid input code can be used in our generic network model, finding activation functions grounded in biological experiments together with matching readout matrices is typically non-trivial. We present one such input model, which resembles fundamental response properties of area MT, in **Section 5.6**.

**Handling $f_\Sigma(\lambda_t^2)$.** The non-linearity of the function $f_\Sigma(\boldsymbol{\lambda}_t^2)$, given by eqn. (42), prohibits a direct incorporation of the effect of $\boldsymbol{\lambda}^2$ on its dependent variables in eqn. (56) and eqn. (57), within the computational framework of **Section 5.4**. The core reason is that the linear readout $\boldsymbol{A}^\lambda \boldsymbol{r}_{\text{dis}}$ does not commute with the square-root function. Yet, it turns out that $f_\Sigma(\boldsymbol{\lambda}_t^2)$ can be incorporated into the network model quite easily owing to its simple functional form. Since $f_\Sigma$ keeps all motion components separate, $f_\Sigma(\lambda_m^2) = \text{const}_m \cdot (-1 + \sqrt{1 + \text{const}_m \lambda_m^2})$, we can employ a dedicated population, $\boldsymbol{r}_{\text{1-to-1}}$, using a one-to-one coding model, which assigns one neuron (or small sub-population) to each component of the posterior variance:

$$r_{\text{1-to-1},m} = \frac{1}{A^\Sigma_{mm}} f_\Sigma(\boldsymbol{A}^\lambda_{m\star} \boldsymbol{r}_{\text{dis}}) \ . \tag{69}$$

Here we have used that $\lambda_m^2 = \boldsymbol{A}^\lambda_{m\star} \boldsymbol{r}_{\text{dis}}$ can be read out linearly from the population's activity. The coefficient $A^\Sigma_{mm}$ scales the firing rate of $r_{\text{1-to-1},m}$. This leads to a neurally plausible activation function of $r_{\text{1-to-1}}$: In **Fig. S9**, replace $\lambda_m^2$ by the "input current" $\boldsymbol{A}^\lambda_{m\star}\boldsymbol{r}_{\text{dis}}$ on the x-axis, and $f_\Sigma$ by $r_{\text{1-to-1}}$ on the y-axis. This is reminiscent of the firing response of leaky integrate-and-fire neurons or, more generally, Type I neurons, as a function of the input current. Finally, we can read out $\boldsymbol{f}_\Sigma$ linearly from $\boldsymbol{r}_{\text{1-to-1}}$ via readout matrix $\boldsymbol{A}^\Sigma = \text{diag}\left[(A^\Sigma_{11}, .., A^\Sigma_{MM})\right]$, thereby allowing us to apply the formalism of **Section 5.4** to $f_\Sigma(\boldsymbol{\lambda}^2)$ (which acts as a variable).

In the above argumentation, we have made two simplifying assumption. First, we have assumed an instantaneous response for $\boldsymbol{r}_{\text{1-to-1}}$ instead of the low-pass filtered response of eqn. (53). Since $f_\Sigma$ varies only on the long time scale $\tau_\lambda$,

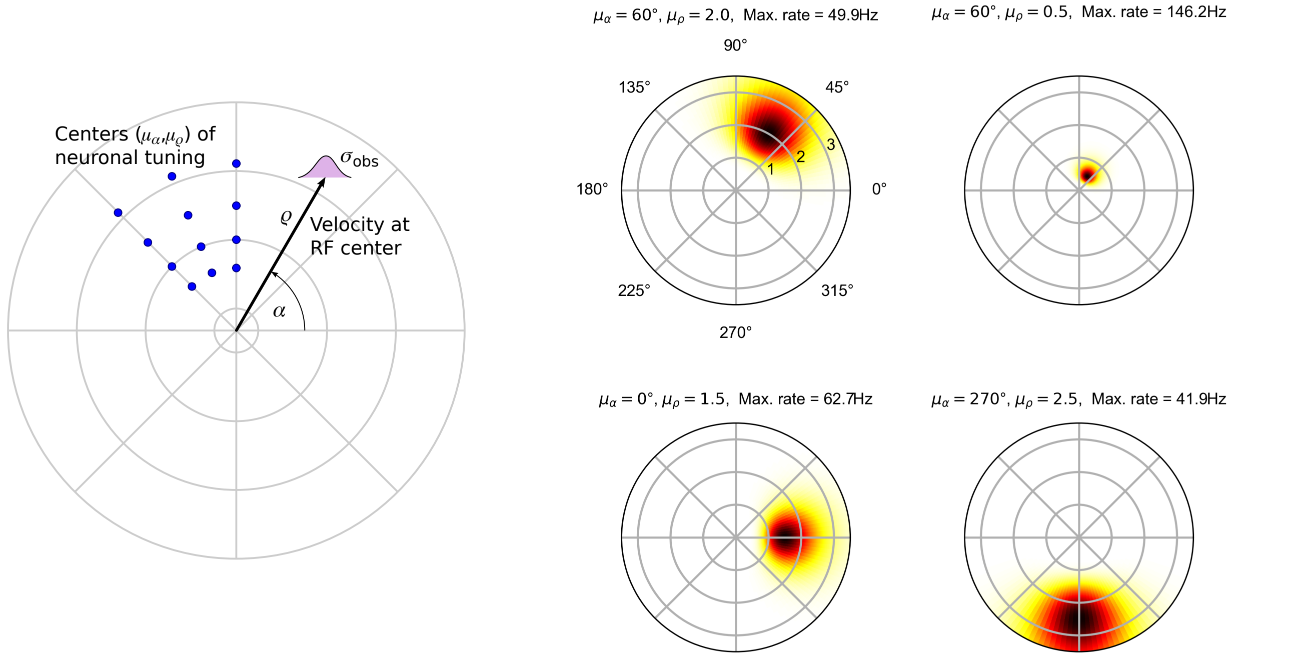

**Figure S11. Tuning functions of MT neurons.** *Left:* The tuning of MT neurons, in response to a stimulus with direction $\alpha$, speed $\rho$ and observation noise $\sigma_{\text{obs}}$, is separable in polar coordinates. In a local population, the neurons' preferred velocity tuning covers directions, $\mu_\alpha$, uniformly, while the density of neurons tuned to speed, $\mu_\rho$, decreases for higher speed. Note that all quantities refer to a local coordinate system centered at the receptive field (RF) center. So, the coordinates here are not to be confused with the coordinate system in **Fig. S8** which describes RF locations. *Right:* Tuning function according to eqn. (70) for four example neurons. Tuning centers and maximum firing rates are given in the axes titles. Parameters (using Python indexing, i.e., $n_\rho = 0, .., N_\rho - 1$): $\psi = 0.1$, $\sigma^2_{\text{obs}} = (0.05/3)^2$, $N_\alpha = 16$, $N_\rho = 12$, $\mu_\rho(n_\rho) = \rho_{\text{min}} + d_\rho\, n_\rho^{1.25}$, $d_\rho = (\rho_{\text{max}} - \rho_{\text{min}})/(N_\rho - 1)^{1.25}$, $\rho_{\text{min}} = 0.1$, $\rho_{\text{max}} = 8.0$, $\kappa_\alpha = 1/0.35^2$, $\sigma^2_\rho = 0.35^2$.

eqn. (69) could easily be replaced by a low-pass filtered version with $f_i := f_\Sigma / A^\Sigma_{mm}$ being the neurons' activation function in eqn. (53). Secondly, we notice that, strictly, $f_\Sigma$ depends not only on $\boldsymbol{\lambda}^2$, but also on $1/\sigma^2_{\text{obs}}$. While the quadratic interaction between these variables, as expressed by eqn. (42), is covered by the theory, we decided to reduce the complexity of the network model by assuming a fixed default value for $\sigma^2_{\text{obs}}$ in the computer simulations. Again, an extension respecting the explicit $\sigma^2_{\text{obs}}$-dependence would be straight-forward.

**The complete network model.** Plugging all of the components together, we obtain the network model shown in **Fig. S10**. This network emulates the adiabatic observer model given by eqn. (43) + (44).

#### 5.6 Neural coding of the input: an example for area MT

While we aimed to leave the neural code for all *latent* variables as generic as possible in the network examples, we specify an *input* code that respects known response properties of area MT. In the following, we present the tuning functions for the input neurons which are derived from models and properties in the literature on area MT (Born and Bradley, 2005; Nover, Anderson, and DeAngelis, 2005; Krekelberg, 2006). Their most important computational property is that they support linear readout of $\boldsymbol{v}/\sigma^2_{\text{obs}}$ and $1/\sigma^2_{\text{obs}}$ in Cartesian coordinates.

We will proceed in three steps. First, we define the tuning functions in polar coordinates since response properties are commonly presented in this coordinate system in the experimental literature. Second, we state some helpful mathematical properties of the proposed tuning functions. Third, we provide the readout matrices $\boldsymbol{A}^v$ and $\boldsymbol{A}^\sigma$ and demonstrate how they accurately decode the relevant variables, $\boldsymbol{v}/\sigma^2_{\text{obs}}$ and $1/\sigma^2_{\text{obs}}$.

**The tuning function in polar coordinates.** Commonly, MT tuning is characterized in polar coordinates because the tuning function becomes separable, i.e., it factorizes into a product of functions over different variables. Owing to the retinotopic organization of MT, we consider for the remainder of this subsection a local population of neurons sharing the same spacial receptive field. Those neurons are indexed, as shown in **Fig. S11 (left)**, by $n_\alpha = 1..N_\alpha$ and $n_\rho = 1..N_\rho$ according to their preferred direction, $\mu_\alpha$, and speed (absolute value of velocity), $\mu_\rho$, respectively. We use the following tuning function in response to a stimulus with direction $\alpha \in [0, 2\pi)$, speed $\rho \geq 0$, and observation noise $\sigma^2_{\text{obs}}$:

$$f(\alpha, \rho, \sigma^2_{\text{obs}}; n_\alpha, n_\rho) = f_\sigma(\sigma^2_{\text{obs}}) \cdot f_\alpha(\alpha; n_\alpha) \cdot f_\rho(\rho; n_\rho) \tag{70}$$

$$\text{with} \qquad f_\sigma(\sigma^2_{\text{obs}}) = \frac{\psi}{\sigma^2_{\text{obs}}} \ , \tag{71}$$

$$f_\alpha(\alpha; n_\alpha) = \frac{d_\alpha}{2\pi I_0(\kappa_\alpha)} \ e^{\kappa_\alpha \cos(\alpha - d_\alpha n_\alpha)} \ , \tag{72}$$

$$f_\rho(\rho; n_\rho) = \frac{\mu'_\rho(n_\rho)}{\sqrt{2\pi\sigma^2_\rho}\,\mu_\rho(n_\rho)} \ e^{-\frac{\left(\log(\rho) - \log(\mu_\rho(n_\rho))\right)^2}{2\sigma^2_\rho}} \ . \tag{73}$$

Example tuning functions are shown in **Fig. S11 (right)**. Eqn. (70) is composed of sub-functions for the noise $f_\sigma$, motion direction $f_\alpha$, and motion speed $f_\rho$, which employ a range of parameters: The overall (maximum) firing rate is scaled by $\psi$. The angle between cells' preferred direction is $d_\alpha = 2\pi/N_\alpha$, such that neuron $n_\alpha$'s preferred direction is $d_\alpha\, n_\alpha$. The directional tuning width is described by $\kappa_\alpha$ (formally, $\kappa_\alpha$ is the precision parameter of a von-Mises-distribution density function, and $I_0(\kappa_\alpha)$ is the modified Bessel function of order 0 for normalization). Neuron $n_\rho$'s preferred speed is given by function $\mu_\rho(n_\rho)$, with $\mu'_\rho$ denoting the function's derivative. Finally, the width of speed tuning is controlled by $\sigma^2_\rho$. Let us briefly discuss how eqn. (70) captures known properties of MT:

- Neurons are tuned to speed (absolute value of velocity) and direction (almost entirely into only one direction, not the opposite direction).
- Direction tuning is commonly described by a von Mises density function. Preferred directions cover the circle roughly isotropically, here via $d_\alpha\, n_\alpha$.
- Speed tuning can be described by a log-normal function of the speed $\rho$. The density of speed tuning centers in MT has been reported to decrease for larger speeds, which can be captured by the function $\mu_\rho(n_\rho)$.
- Activity is modulated by contrast (via $\sigma^2_{\text{obs}}$), with lower contrast (higher noise) attenuating the overall firing rate.

We make the simplifying assumptions that (i) all neurons have the same firing rate scaling factor $\psi$, and (ii) that the tuning widths, given by $\kappa_\alpha$ and $\sigma^2_\rho$, are "not too wide". The meaning of "not too wide" will become clear in the following mathematical consideration.

**Mathematical properties of the tuning function.** We next discuss some useful properties of the components of the above tuning function. First, we note that $f_\alpha$ and $f_\rho$ have the form of known probability density functions over the neuron indices $n_\alpha$ and $n_\rho$, respectively. In particular, they integrate to one in the limit of many, narrowly spaced neurons:

$$\int_0^{\frac{2\pi}{d_\alpha}} f_\alpha(\alpha; n_\alpha)\, dn_\alpha = \int_0^{2\pi} e^{\kappa_\alpha \cos(\alpha - d_\alpha n_\alpha)} / (2\pi I_0(\kappa_\alpha))\, d(d_\alpha n_\alpha) = 1 \tag{74}$$

$$\text{and} \qquad \int_0^\infty f_\rho(\rho; n_\rho)\, dn_\rho = \int_0^\infty \frac{1}{\sqrt{2\pi\sigma^2_\rho}\,\mu_\rho} e^{-\frac{\left(\log(\rho) - \log(\mu_\rho)\right)^2}{2\sigma^2_\rho}} \ d\mu_\rho = 1 \ . \tag{75}$$

Furthermore, the distributional forms give rise to nice moments w.r.t. the tuning centers $d_\alpha\, n_\alpha$ and $\mu_\rho(n_\rho)$:

$$\langle e^{i\, d_\alpha n_\alpha} \rangle_{f_\alpha} = \frac{I_1(\kappa_\alpha)}{I_0(\kappa_\alpha)} e^{i\alpha} \overset{\text{large}\ \kappa_\alpha}{\approx} e^{i\alpha} = \begin{pmatrix} \cos\alpha \\ \sin\alpha \end{pmatrix} \tag{76}$$

$$\text{and} \qquad \langle \mu_\rho(n_\rho) \rangle_{f_\rho} = \rho e^{\sigma^2_\rho/2} \overset{\text{small}\ \sigma_\rho}{\approx} \rho \ . \tag{77}$$

We now understand how narrow (i.e., "not too wide") tuning functions enable reading out the encoded direction, $\alpha$, and speed, $\rho$: large $\kappa_\alpha$ and small $\sigma^2_\rho$ afford the approximations in eqn. (76) and (77). Further, the mathematical relations highlight that $\kappa_\alpha$ and $\sigma^2_\rho$ could be modulated by the observation noise $\sigma^2_{\text{obs}}$ without changing the ability to encode/decode the input.

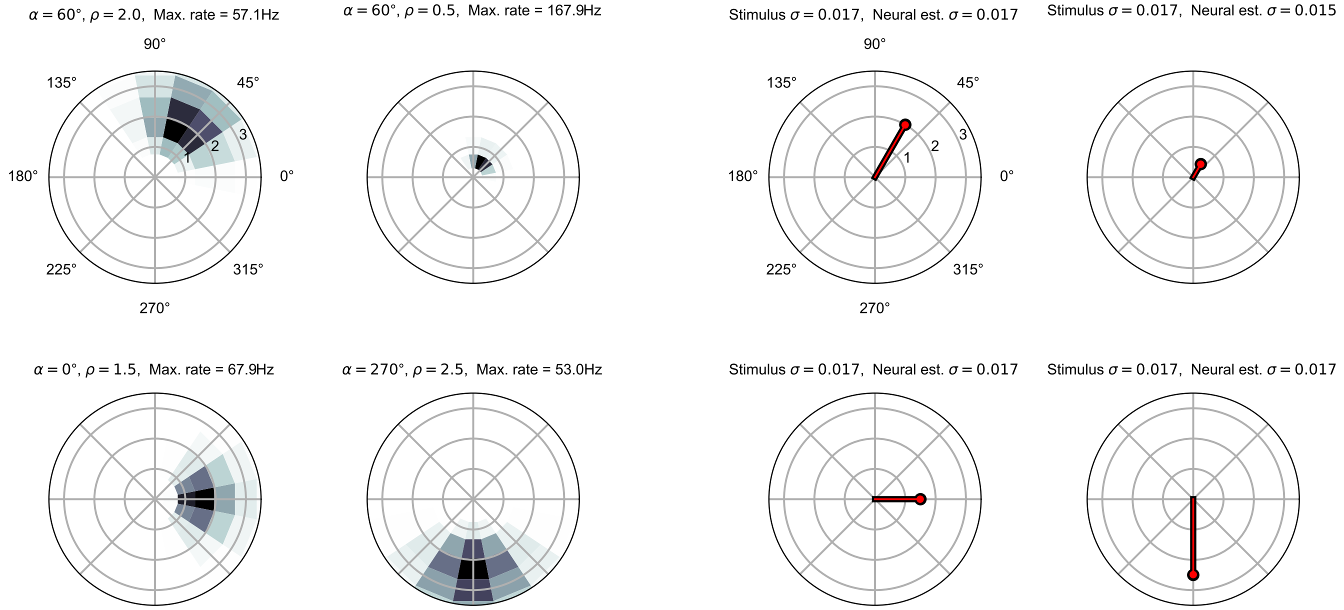

**Figure S12. Linear readout of input statistics from population responses.** *Left:* Population response when encoding four example stimuli. The population consists of 192 neurons with the parameters given in **Fig. S11**. For clarity, only neurons with $\mu_\rho < 3.5$ are shown. *Right:* Linear readout of $1/\sigma^2_{\text{obs}}$, $v_x/\sigma^2_{\text{obs}}$ and $v_y/\sigma^2_{\text{obs}}$ from the population activities on the left via weights given by eqn. (78). Shown are, in polar coordinates, the stimulus ground truth (black) and the estimate by the linear readout (red). The estimated uncertainty is provided is the axes titles.

**Linear readout of input statistics.** With the above mathematical properties at hand, we identify matrices $\boldsymbol{A}^\sigma$ and $\boldsymbol{A}^v$ for linear readout:

$$A^\sigma_{k,(n_\alpha,n_\rho)} = \frac{1}{\psi} \qquad \text{and} \qquad A^v_{k,(n_\alpha,n_\rho)} = \frac{1}{\psi}\begin{pmatrix}\cos d_\alpha n_\alpha \\ \sin d_\alpha n_\alpha\end{pmatrix}\mu_\rho(n_\rho)\ , \tag{78}$$

because reading out with these matrices from the MT-population of the $k$-th observable yields:

$$\int\!\!\int A^\sigma_{k,(n_\alpha,n_\rho)}\, f(\alpha_k,\rho_k,\sigma^2_{\text{obs},k};\, n_\alpha, n_\rho)\, dn_\alpha\, dn_\rho = \frac{1}{\sigma^2_{\text{obs},k}} \tag{79}$$

$$\int\!\!\int A^v_{k,(n_\alpha,n_\rho)}\, f(\alpha_k,\rho_k,\sigma^2_{\text{obs},k};\, n_\alpha, n_\rho)\, dn_\alpha\, dn_\rho = \frac{\rho_k}{\sigma^2_{\text{obs},k}}\begin{pmatrix}\cos\alpha_k \\ \sin\alpha_k\end{pmatrix} = \begin{pmatrix}v_x/\sigma^2_{\text{obs},k} \\ v_y/\sigma^2_{\text{obs},k}\end{pmatrix}\ . \tag{80}$$

Examples of the population response to four motion stimuli is shown in **Fig. S12 (left)** for a population of 192 neurons. The tuning centers span 12 radii ("speed") and 16 angles ("direction"). In **Fig. S12 (right)**, the resulting linear readout (red) is shown alongside the ground truth (black). Even the coarse coverage of the stimulus space by 192 neurons is sufficient for a faithful reconstruction of the stimulus. As an interesting observation, while neurons tuned to higher speeds have wider tuning curves, this does not imply that their activity would encode heightened uncertainty: $\sigma_{\text{obs}}$ is identical in all of the examples in **Fig. S11** and **Fig. S12**.

In summary, we have identified with eqn. (70) an MT tuning function that supports linear readout of the variables $\boldsymbol{v}/\sigma^2_{\text{obs}}$ and $1/\sigma^2_{\text{obs}}$ via matrices $\boldsymbol{A}^v$ and $\boldsymbol{A}^\sigma$.

#### 6. Captions to the supplemental videos

**Video S1: Johansson experiment.** *Top:* 3-dot motion display. *Bottom left:* Evolution of motion strengths, $\lambda_m(t)$. *Bottom right:* Motion sources, $s_m(t)$. The stimulus is decomposed into horizontal shared motion (magenta) and vertical individual motion (green) for the central dot.

**Video S2: Duncker wheel.** *Top:* Duncker wheel motion display. *Bottom left:* Evolution of motion strengths, $\lambda_m(t)$. *Bottom right:* Motion sources, $s_m(t)$. The stimulus is decomposed into rightward shared motion (magenta) and clockwise rotation (green) for the dot on the rim.

**Video S3: Motion illusion from Lorenceau (1996).** Motion displays without and with motion noise are presented. Multiple noise levels are used because the noise threshold for inducing the perceptual switch is participant-dependent. To prevent priming on a certain grouping, different random groups were drawn for every noise level, in this demo video. (The simulation in Fig. 4 used, apart from $\sigma^2_{obs}$, identical inputs.)

**Video S4: Nested structure-from-motion displays.** A series of videos covering all SfM conditions from Figure 6. The "normal" rotation speed is 90 °/s, the "faster" speed is 135 °/s. The outer cylinder is 50% larger than the inner cylinder, by radius. In the videos, points have an average life time of 250 ms to preclude object-indexed tracking of dots.

**Video S5: Location-indexed stimulus with varying fraction of shared motion.** Every 10 s, the fraction of shared motion across the apertures increases by 1/8, ranging from $q=1/8$ (almost independent motion) to $q=7/8$ (almost fully correlated motion). Steps in $q$ are marked by brief flashes of gray squares.

Nover, Harris, Charles H. Anderson, and Gregory C. DeAngelis (Oct. 2005). "A Logarithmic, Scale-Invariant Representation of Speed in Macaque Middle Temporal Area Accounts for Speed Discrimination Performance". In: *Journal of Neuroscience*. Publisher: Society for Neuroscience Section: Behavioral/Systems/Cognitive.

Salinas, Emilio and Laurence F Abbott (1996). "A model of multiplicative neural responses in parietal cortex". In: *Proceedings of the national academy of sciences*.

Weiss, Yair, Eero P Simoncelli, and Edward H Adelson (2002). "Motion illusions as optimal percepts". In: *Nature neuroscience*.
